# Mixed Effects Models for Resampled Network Statistics Improves Statistical Power to Find Differences in Multi-Subject Functional Connectivity

**DOI:** 10.1101/027516

**Authors:** Manjari Narayan, Genevera I. Allen

**Affiliations:** Department of Electrical and Computer Engineering; Department of Statistics, Rice University, Houston, TX, USA; Jan and Dan Duncan Neurological Research Institute, Houston, TX, USA

**Author notes:** Correspondence*: Manjari Narayan Department of Electrical and Computer Engineering, Rice University, 6100 Main St, Houston, TX, 77005, USA.

## Abstract

Many complex brain disorders, such as autism spectrum disorders, exhibit a wide range of symptoms and disability. To understand how brain communication is impaired in such conditions, functional connectivity studies seek to understand individual differences in brain network structure in terms of covariates that measure symptom severity. In practice, however, functional connectivity is not observed but estimated from complex and noisy neural activity measurements. Imperfect subject network estimates can compromise subsequent efforts to detect covariate effects on network structure. We address this problem in the case of Gaussian graphical models of functional connectivity, by proposing novel two-level models that treat both subject level networks and population level covariate effects as unknown parameters. To account for imperfectly estimated subject level networks when fitting these models, we propose two related approaches — R^2^ based on resampling and random effects test statistics, and R^3^ that additionally employs random adaptive penalization. Simulation studies using realistic graph structures reveal that R^2^ and R^3^ have superior statistical power to detect covariate effects compared to existing approaches, particularly when the number of within subject observations is comparable to the size of subject networks. Using our novel models and methods to study parts of the ABIDE dataset, we find evidence of hypoconnectivity associated with symptom severity in autism spectrum disorders, in frontoparietal and limbic systems as well as in anterior and posterior cingulate cortices.

## 1 INTRODUCTION

One of the goals of neuroimaging studies of intrinsic or “resting state” brain activity, is to discover specific and stable imaging based biomarkers or phenotypes of neuropsychiatric and neurological disorders. Typically, resting state studies seek to infer *functional connectivity* or functional relationships between distinct brain regions from observed neurophysiological activity. Advances in resting state studies using fMRI [**Bullmore**, 2012; **Menon**, 2011; **Craddock et al.**, 2013; **Smith et al.**, 2013] suggest that functional connectivity could yield neuroimaging biomarkers for the diagnosis and personalized treatment of a wide range of disorders.

For instance, many studies have found differences either in individual functional connections or in overall patterns of connectivity in autism spectrum disorders [**Uddin**, 2014; **Di Martino et al.**, 2014a], Alzheimer’s [**Buckner et al.**, 2009; **Tam et al.**, 2014], major depression [**Tao et al.**, 2013; **Lui et al.**, 2011; **Kaiser et al.**, 2015] and others [**Meda et al.**, 2012; **van den Heuvel et al.**, 2013; **Palaniyappan et al.**, 2013]. However, simple group level differences between two distinct samples are challenging to interpret in many disorders. Autism, for example, is a diagnostic label that masks many diverse clinical symptoms [**Lenroot and Yeung**, 2013; **Insel**, 2014]. Thus, the biological relevance of group level differences in network structure between autism and healthy populations is unclear for individual subjects. One solution to find more meaningful differences in network structure is to study whether behavioral and affective symptoms measured by cognitive scores are associated with variations in individual functional networks. This paper offers a novel and rigorous statistical framework to find and test such covariate effects on functional connectivity metrics, when functional connectivity is defined using Gaussian graphical models.

Functional connectivity refers to latent relationships that cannot be directly observed via any modality of functional neuroimaging. Instead, it must be estimated from observations of neurophysiological activity. In fMRI studies, we first observe changes in the BOLD response over time either across thousands of voxels or over hundreds of brain regions, defined anatomically or functionally. Then depending on the specific statistical definition for functional connectivity, we estimate a functional connectivity network per subject using within-subject BOLD observations. For example, in a pairwise correlation model of functional connectivity, if the mean time-series of two brain regions are correlated then they are functionally connected. Thus, one popular approach to estimate functional connectivity is to compute sample correlations between every pair of brain regions. An increasingly popular alternative is to use Gaussian graphical models (GGMs) based on partial correlations to define functional connectivity. Here, if two brain regions are partially correlated, that is if the mean time-series of two brain regions remain correlated after regressing out the time-series of other brain regions, then they are functionally connected. For multivariate normal data, a zero partial correlation between two brain regions is equivalent to independence between the activity of two brain regions conditional on the activity of all other intermediate brain regions. Thus, GGMs eliminate indirect connections between regions provided by pairwise correlations and are increasingly popular in neuroimaging [**Marrelec et al.**, 2006; **Smith et al.**, 2011; **Varoquaux et al.**, 2012; **Craddock et al.**, 2013]. Consequently, employing GGMs for functional connectivity enables us to discover network differences that implicate nodes and edges directly involved in producing clinical symptoms, and provide stronger insights into network structures truly involved in the disease mechanism. For the rest of this paper, we define functional connectivity in terms of GGMs and discuss approaches to conducting inference on network metrics for such network models.

The functional connectivity of a single experimental unit or subject is rarely the final object of interest. Rather, most neuroimaging studies [**Bullmore and Sporns**, 2012; **Zuo et al.**, 2012; **Bullmore**, 2012] are interested in identifying network biomarkers, or broader patterns of functional connectivity shared across individuals who belong to some distinct population or display some clinical phenotype. A popular approach [**Bullmore and Sporns**, 2009] to find such network biomarkers is through topological properties of network structure. Common properties or metrics either measure specialization of network components into functionally homogenous modules, or measure how influential brain regions integrate information across distinct network components. However, recall that functional connectivity in individual subjects is unknown and unobserved. Consequently, many multi-subject fcMRI studies first estimate functional connectivity for every subject and then assuming these subject networks are fixed and known, compute topological metrics of these networks using the Brain Connectivity Toolbox [**Rubinov and Sporns**, 2010]. Finally, they compare and contrast these estimated networks or estimated network metrics to infer group level network characteristics. Typical neuroimaging studies that seek to detect covariate effects on network structure [**Hahamy et al.**, 2015; **Warren et al.**, 2014] conduct a single level regression with network metrics as the response and cognitive scores as the covariate, and subsequently use standard F-tests for covariate testing. New methods to conduct such network inference either emphasize novel topological metrics [**van den Heuvel and Sporns**, 2011; **Alexander-Bloch et al.**, 2012] or novel approaches to study covariate effects for known networks for complex experimental designs with longitudinal observations or multiple experimental conditions [**Simpson et al.**, 2013; **Kim et al.**, 2014; **Ginestet et al.**, 2014]. However, these existing approaches assume estimated functional networks are perfectly known quantities. In contrast, we seek to explicitly investigate the consequences of using estimated, and often imperfectly estimated, functional networks and their corresponding network metrics on subsequent inference for covariate effects.

Before considering the consequences of using estimated networks, one might ask the question why individual network estimates might be unreliable to begin with. Statistical theory informs us that estimated networks can be unreliable in two possible ways. First, high dimensional networks with a large number of nodes estimated from a limited number of fMRI observations in a session possess substantial sampling variability [**Narayan et al.**, 2015; **Bickel and Levina**, 2008; **Rothman et al.**, 2008; **Ravikumar et al.**, 2011]. Second, when assuming sparsity in the network structure in the form of thresholded or penalized network estimates to overcome high dimensionality, we often obtain biased network estimates in the form of false positive or false negative edges [**Ravikumar et al.**, 2011]. Such errors in estimating networks are particularly exacerbated [**Narayan et al.**, 2015] when networks are well connected with modest degrees, as is the case in neuroimaging. Additionally, empirical evidence from neuroimaging studies also suggest that sample correlation based estimates of individual resting state networks are unreliable. For instance test re-test studies [**Shehzad et al.**, 2009; **Van Dijk et al.**, 2010; **Braun et al.**, 2012] that measure intersession agreement of estimated functional networks within the same subject find that sample intra-class correlations vary between.3 – .7, indicating non-negligible within subject variability. While we expect many sources of variation contribute to such inter-session variability within a subject including natural variations due to differences in internal cognitive states, recent work by **Birn et al.**, [2013]; **Hacker et al.**, [2013]; **Laumann et al.**, [2015] suggests that sampling variability due to limited fMRI measurements play a significant role. These studies find that increasing the length of typical fMRI sessions from 5-10 minutes to 25 minutes substantially improves inter-session agreement of functional networks. Given the accumulating theoretical and empirical evidence of these methodological limitations, we assume that obtaining perfect estimates of individual networks is unlikely in typical fMRI studies. Instead, we seek to highlight the importance of accounting for imperfect estimates of functional networks in subsequent inferential analyses.

Failure to account for errors in estimating statistical networks reduces both generalizability and reproducibility of functional connectivity studies. Statistical tests that compare functional networks but do not account for potentially unreliable network estimates lack either statistical power or type I error control or both. For instance, **Narayan and Allen** [2013]; **Narayan et al.**, [2015] investigate the impact of using estimated networks when testing for two-sample differences in edge presence or absence between groups. When individual subject graphical models cannot be estimated perfectly, **Narayan et al.**, [2015] show that standard two-sample test statistics are both biased and overoptimistic, resulting in poor statistical power and type I error control. Though this paper is similar in spirit to previous work [**Narayan et al.**, 2015] in emphasizing the adverse effects of using estimated networks to study differences in functional connectivity, the unique contribution of this work are as follows: (1) Whereas previous work considered simple two-sample tests, we consider general covariate effects (that include both binary and continuous covariates) to link symptom severity to individual variations in functional connectivity. (2) We propose methods relevant to network metrics beyond the edge level. Finally, we provide empirical results such as statistical power analyses that offer greater practical guidance on choosing sample size and planning data analysis for future studies.

The paper is organized as follows. In Section 2 we provide new statistical models that explicitly link subject level neurophysiological data to population level covariate effects for network metrics of interest and provide new statistical algorithms and test statistics using resampling and random penalization for testing covariate effects. While the models and methods we propose can detect covariate effects on many well behaved network metrics [**Balachandran et al.**, 2013] at the edge level [**Tomson et al.**, 2013], node level [**Buckner et al.**, 2009; **Zuo et al.**, 2012] and community level [**Tomson et al.**, 2013; **Alexander-Bloch et al.**, 2012], we investigate the benefits of our methods to discover covariate effects on connection density. Using realistic simulations of graph structure for GGMs in Section 3, we demonstrate our proposed resampling framework substantially improves statistical power over existing approaches, particularly for typical sample size regimes in fMRI studies. Finally, in Section 4 we demonstrate that our proposed methods can detect biologically relevant signals in a resting state fMRI dataset for autism spectrum disorders.

## 2 MODELS AND METHODS

We seek new methods to detect covariate effects when populations of functional networks are unknown. To achieve this, we first need statistical models that describe how each measurement of brain activity denoted by 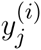 arises from unknown functional brain network with *p* nodes in the *i*^th^ subject and how individual variations in a population of brain networks are related to some population level mean. Thus, for any network model and any network metric under investigation, we propose the following general two-level models to investigate covariate effects in functional connectivity. In subsequent sections, we provide specific instances of these models investigated in this paper.

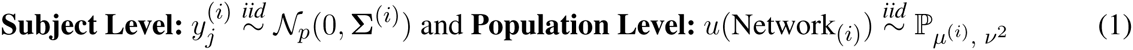

where Σ^(*i*)^ is the covariance, Network_*i*_ is an adjacency matrix derived from either the covariance, the inverse covariance Θ = (Σ^(*i*)^)^−1^ or their correlational counterparts and *u*(·) denotes some network metric over the brain network. In this paper, we assume the individual measurements of brain activity at the subject level follow a multivariate normal distribution. At the population level, we assume that the effect of covariates on the network metrics follows a generalized linear model [**Searle et al.**, 2009] where the mean and variance of the relevant continuous or discrete probability distribution, ℙ, for the network metric of interest is given by *μ*^(*i*)^ and *ν*^2^.

Suppose that we denote any network metric in the *i*^*th*^ subject as *u*^(*i*)^ and the vector of network metrics as u = (*u*^(1)^,…, *u*^(*n*)^), then the population mean is given by *μ* = **𝔼**(u) and population variance is given by Var(*u*^(*i*)^ = *ν*^2^. Then the generalized linear model for the population mean is given by

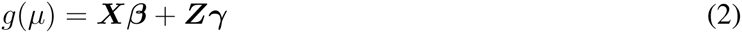

Here *g*(*μ*) is a link function either reduces to *g*(*μ*) = *μ* in linear models, or takes other forms such as the logit functon for nonlinear models; ***X*** is the *n* × (*q* + 1) matrix of the intercept and *q* covariates of interest with corresponding coefficients ***β*** = (*β*_0_, *β*_1_,… *β*_*q*_) while *Z* is the *n* × *r* matrix of nuisance covariates and corresponding regression coefficients γ. *X*_*i*_ and *Z*_*i*_ denote the *q* dimensional explanatory covariate and *r* dimensional nuisance covariate for the *i*^*th*^ subject, respectively.

In this paper, we seek to test the hypothesis that explanatory covariates have a statistically significant covariate effect on network metrics. Here *β*_\0_ denotes the coefficients for explanatory covariates. Thus, the null ***H***_0_ and alternative hypothesis ***H***_1_ are

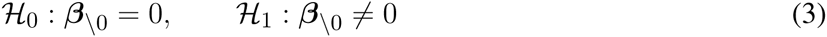

This section is organized as follows — In Section 2.1, we specifically employ Gaussian graphical model of functional connectivity at the subject level and investigate covariate effects using linear models for density based network metrics for the population level. Standard statistical analyses in neuroimaging studies estimate each level of these two level models separately. Thus, such approaches first estimate functional connectivity networks by fitting subject level models. However, they assume individual subject networks and their metrics are known when they fit the population level model and conduct inference on covariate effects. In Section 2.2 we discuss how such statistical procedures that assume functional connectivity networks are known lose statistical power to detect covariate effects. To address this problem, we introduce two related methods that utilize resampling, random adaptive penalization, and random effects that we call, **R**^2^ and **R**^3^ in Section 2.3. These methods ameliorate potential biases and sampling variability in estimated network metrics, thus improving statistical power to detect covariate effects.

### 2.1 TWO LEVEL MODELS FOR COVARIATE EFFECTS

We begin by studying the earlier subject level network model in (1) specifically for networks given by Gaussian graphical models. Recall that the *p*-variate random vector 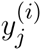 denotes BOLD observations or average BOLD observations within *p* regions of interest, at the *j*^*th*^ time point for the *i*^*th*^ subject. We assume 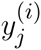 has a multivariate normal distribution,

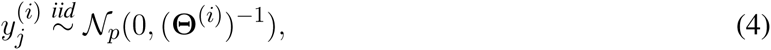

where the network model of interest is derived from the inverse covariance or precision matrix Θ^(*i*)^, *j* = 1,… *t*, and *i* = 1,… *n*. In subsequent sections, we denote the *t* × *p* data matrix of observations by 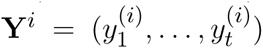 and the random variable associated with each brain region as *Y*_*k*_. Although fMRI observations are autocorrelated across time and thus dependent [**Worsley et al.**, 2002; **Woolrich et al.**, 2001], we assume that these observations can be made approximately independent via appropriate whitening procedures discussed in our case study in Section 4.

Let ***G*** (*ν*, *ε*) denote a Gaussian graphical model that consists of vertices *ν* = {1, 2,…,*p*} and edges *ε* ⊂ *ν* × *ν*. Here, the presence of an edge (*k*,*l*) ϵ *ε* implies that the random variables *Y*_*k*_ and *Y*_*l*_ at nodes/vertices *k* and *l* are statistically dependent conditional on all the other vertices *ν* \ {*k*, *l*}. For multivariate normal distributions, a non-zero value in the (*k*, *l*) entry of the inverse covariance matrix Θ^(*i*)^ is equivalent to the conditional independence relationships, *Y*_*k*_ ⊥ *Y*_*l*_ |*Y*_*V*\{*k*,*l*}_. Thus, we define functional connectivity networks where edges indicate *direct* relationships between two brain regions using the non-zero entries of Θ^(*i*)^. For a more thorough introduction to graphical models, we refer the reader to **Lauritzen** [1996].

Following the neuroimaging literature [**Bullmore and Sporns**, 2009], we consider network metrics to be functions of a binary adjacency matrix. The adjacency matrix of each individual subject network in our model (4) is given by the support of the inverse covariance matrix 𝕀{Θ^(*i*)^ ≠ 0}. Network metrics that measure topological structure of networks are widely used in neuroimaging [**Bullmore and Sporns**, 2009; **Rubinov and Sporns**, 2010]. While any of these network metrics can be incorporated into our two level models, we have found that many metrics originally proposed when studying a determinstic network are not suitable for covariate testing in the presence of individual variations in a population of networks. Recently, **Balachandran et al.**, [2013] suggests that several discontinuous network metrics which include betweenness centrality, clustering coefficients defined at the node level and potentially many others are not suitable for inference. Thus, this paper focuses on well behaved topological metrics, namely density based metrics. Formally, the density or number of connections in any binary adjacency matrix *A* is given 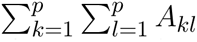. However, rather than defining density over the whole graph, the density can be restricted to a subnetwork (subnetwork density) or over a single node (node density or degree) or simply at the edge level (edge presence). At the node level, density is a simple measure of influence or centrality of a single brain region of interest [**Rubinov and Sporns**, 2010; **Power et al.**, 2013]. At the subnetwork level, density is popularly used [**Honey et al.**, 2007; **Bullmore and Sporns**, 2009] to measure an excess or deficit of long range connections either within or between groups of brain regions with a distinct functional purpose. While we investigate node and subnetwork density in this paper, alternative network metrics amenable to inference include binary metrics such as edge presence [**Narayan et al.**, 2015; **Meda et al.**, 2012] or co-modularity relationships between nodes [**Tomson et al.**, 2013; **Bassett et al.**, 2013].

#### 2.1.1 Population Model for Network Metrics

As described earlier, given the subject level model and a network metric of interest, we use a generalized linear model in (2) to describe the deterministic relationship between the population mean for the network metrics and various covariates of interest. Depending on whether a network metric is continuous or binary valued, this general linear model takes the form of linear or logistic-linear models.

However, we also require a probability model to describe how a random sample of individual network metrics deviate from the population mean. When the network metric *u*^(*i*)^ is continuous valued, the link function in (2) reduces to the identity *g*(*μ*) = *μ*. For network metrics *u*^(*i*)^ such as global, subnetwork or node density, we use the following linear model with normal errors,

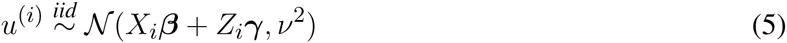

For metrics such as edge presence and co-modularity that take discrete binary values {0,1}, a widely used link function [**Agresti**, 2002; **Williams**, 1982] for the generalized linear model (2) is the logit function. The resulting logistic-linear model takes the following form

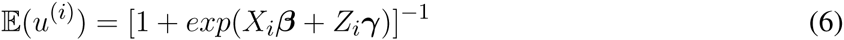

For the remainder of this paper, we consider normal models for node and subnetwork density.

### 2.2 MOTIVATION FOR NEW TEST STATISTICS

To understand why new statistical methods are necessary to fit our two-level models, consider the our covariate testing problem (3) for node and subnetwork density. Suppose the subject level networks in (4) and corresponding metrics are known precisely for each subject. In this case, we employ standard least squares estimation with corresponding F-tests for linear regression to test our null hypothesis for covariate effects (3).

In practice however, not only is the covariate effect *β* unknown, the underlying graphical model Θ^(*i*)^ and the network metric *u*^(*i*)^ is also unknown and are all estimated from data. In Figures 1a and 1b we contrast the ideal scenario where the population of networks and corresponding network metrics are exactly known with the practical scenario where these network metrics are estimated from data. (See Section 3.1 for details on how we simulate data.) Applying a standard linear regression to known network metrics reveals an oracle estimate of the covariate effect (blue line). In contrast, when the standard approach described is applied to estimated network metrics (orange line), the size of the covariate effect is substantially reduced. However, by employing the **R**^3^ approach (green line) that we introduce in the next section, we account for errors in estimating networks, thereby improving statistical power.

**Figure 1.**
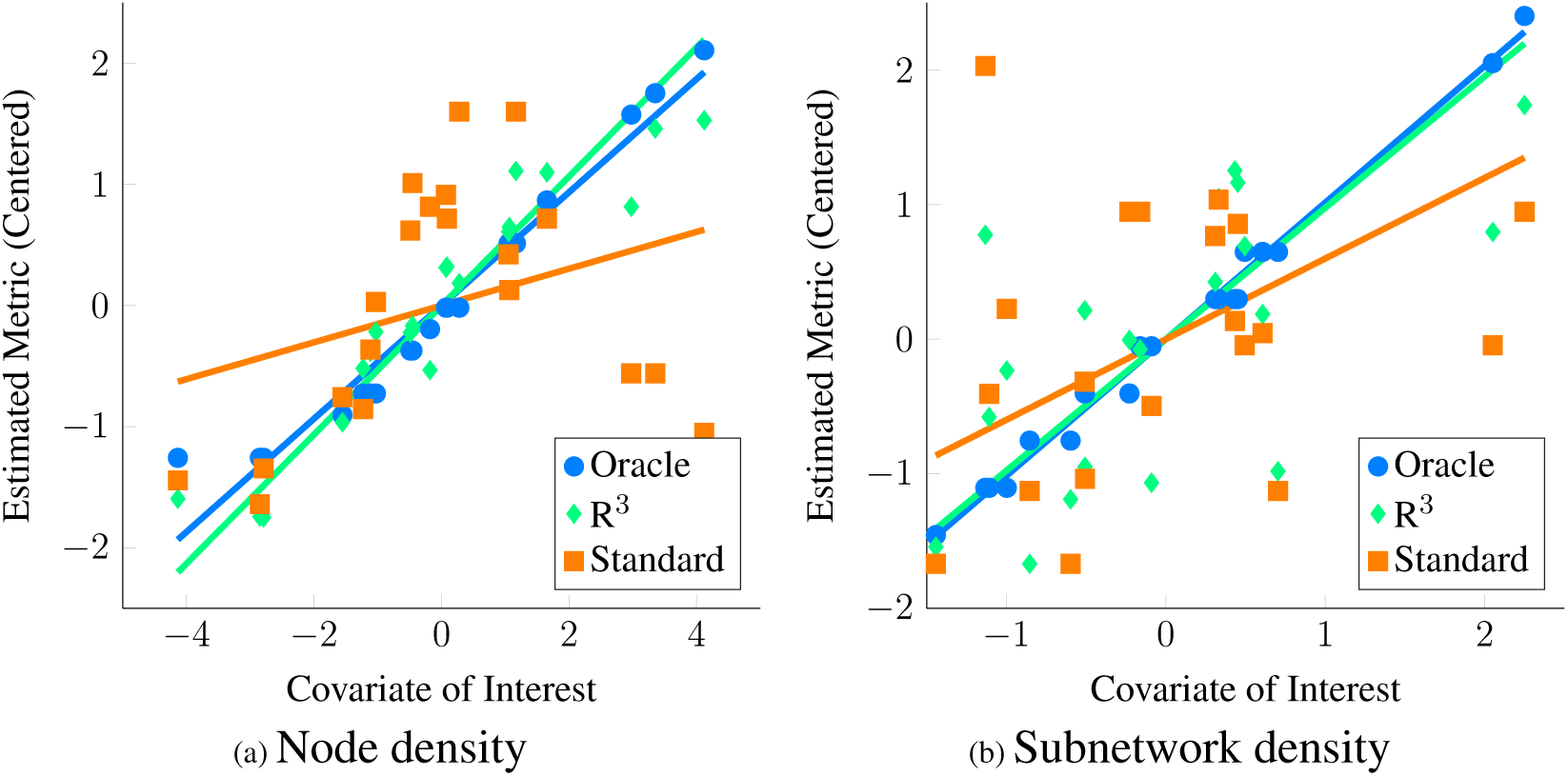
Motivation for new statistical framework **R**^3^. Here, we simulate covariate effects on the metric of interest, namely the degree centrality or node density (left) and subnetwork density (right) with (*p* = 50, *n* = 20, *t* = 200). We illustrate covariate effects in the ideal scenario where network metrics are known perfectly in blue. Unfortunately, in functional connectivity networks, statistical errors in estimating graphical models are inevitable and these propagate to estimates of network metrics. As a result, when we estimate node and subnetwork density for each subject and conduct tests for covariate effects using standard F-tests, we fail to see a clear relationship between metrics and covariate of interest (orange) using linear regression. This loss of statistical power occurs when standard test statistics assume that estimates of density are correct. In contrast, when we account for errors in graph estimation and selection using **R**^3^ test statistics (green), we have greater statistical power to detect covariate effects on density metrics. Algorithmic details of the **R**^3^ approaches can be found in Section 2.

Two issues arise when we estimate network metrics from data. First, instead of true network metrics, *u*^(*i*)^, our estimated network metrics, *ũ*^(*i*)^, are a function of observations **Y**^(*i*)^. Thus, each estimate, *ũ*^(*i*)^, possesses additional sampling variability. However, since we only acquire one network estimate per subject, standard least squares estimation cannot account for this additional variability. Additionally graph selection errors in network estimation potentially bias network metric estimates. Previously, **Meinshausen and Biihlmann** [2006]; **Ravikumar et al.**, [2011]; **Narayan et al.**, [2015] show that in finite sample settings where the number of independent observations *t* within a subject is comparable to the number of nodes *p*, we expect false positive and false negative edges in network estimates. Such graph selection errors increase with the complexity of the network structure, governed by factors such as the level of sparsity, maximum node degree as well as the location of edges in the network. Since functional connectivity networks are moderately dense and well connected with small world structure [**Achard et al.**, 2006], edges in these networks might be selected incorrectly. Observe that in Figures 1a and 1b, we obtain larger estimates of node and subnetwork density for individual networks where true node or subnetwork densities are small and the reverse for truly large node or subnetwork densities. As a result, individual variation in estimated metrics no longer reflects the true effect of the covariate, resulting in loss of statistical power. For a detailed overview of how selection errors in estimating network structure propagate to group level inferences, we refer the reader to Section 2 of **Narayan et al.**, [2015].

To overcome these obstacles, we use resampling to empirically obtain the sampling variability of estimated network metrics, *ũ*^(*i*)^, and propagate this uncertainty using mixed effects test statistics for the covariate effect 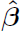. Moreover, by aggregating network statistics across resamples and optionally incorporating adaptive penalization techniques, we sufficiently improve network estimates and corresponding network metrics to obtain more accurate estimates of the covariate effects.

### 2.3 PROCEDURE FOR TESTING COVARIATE EFFECTS

In order to improve statistical power, we propose a resampling framework that integrates network estimation with inference for fixed covariate effects at the population level. We provide two related procedures to test covariate effects – **R**^2^ that employs resampling (RS) and random effects test statistics (RE), and **R**^3^ that employs resampling (RS), random adaptive penalization (RAP) and random effect test statistics (RE). Intuitively, our algorithm consists of first obtaining initial estimates of the sparsity levels in individual subject networks. Then, to estimate the sampling variability of each subject network empirically, we resample within subject observations and re-estimate the networks of each subject. Additionally, in the case of **R**^3^ we simultaneously apply random adaptive penalties when re-estimating the networks. Network metrics are computed on each of the resampled networks, giving us multiple pseudoreplicates of network metrics per subject. Finally, we model these resampled network statistics using simple mixed effects models to derive test statistics for population level covariate effects. After performing our procedure, one can use well known parametric or non-parametric approaches to obtain *p*-values and correct for multiplicity of test statistics when necessary. Thus our resampling framework consists of three components, graph estimation and selection, resampling and optionally RAP, and covariate testing via mixed effects models. We discuss each of these ingredients separately before putting them together in Algorithm 1.

#### 2.3.1 Graphical Model Estimation

Many approaches such as sparse regularized regression [**Meinshausen and Biihlmann**, 2006], sparse penalized maximum likelihood (ML) or the graphical lasso [**Yuan and Lin**, 2007; **Friedman et al.**, 2008] and others [**Cai et al.**, 2011; **Zhou et al.**, 2011] can be used to estimate Θ^(*i*)^ in our subject level model (4). We use the QuIC solver [**Hsieh et al.**, 2011, 2013] to fit a weighted graphical lasso to obtain estimates of Θ^(*i*)^.

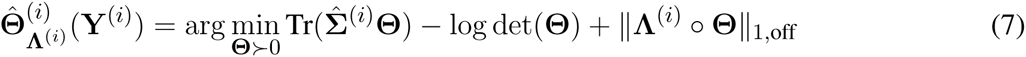

where 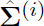 is the empirical sample covariance, 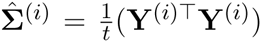, and `o` denotes the Hadamard dot product. The term ||Θ||_1,off_ = ∑_*k*<*l*_ |*θk*,*ι*| is the *ℓ*_1_ penalty on the off-diagonals entries. Since the sample correlation rather than covariance is commonly used in neuroimaging, we employ sample correlation matrix, Σ̃^(*i*)^. The two are equivalent when **Y**^(*i*)^ has been centered and scaled. Given any estimate of the inverse covariance matrix 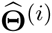, the estimated adjacency matrix for each subject is thus given by 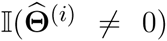 and network statistics can be computed accordingly. For our **R**^3^ procedure, we employ a symmetric weight matrix of penalties **∧**^(*i*)^ obtained by randomly perturbing an initial penalty parameter *λ*^(*i*)^. For our **R**^2^ this weight matrix **∧**^(*i*)^ reduces to a scalar value *λ*^(*i*)^ for all off-diagonal entries, giving us the standard graphical lasso. In order to estimate these initial penalty parameters *λ*^(*i*)^, we employ StARS [**Liu et al.**, 2010], a model selection criterion that is asymptotically guaranteed to contain the true network, and works well with neuroimaging data. The *beta* parameter of StARS is set to.1 in our work.

#### 2.3.2 Resampling and Random Adaptive Penalization

Since network estimates depend on the underlying observations **Y**^(*i*)^, we employ resampling techniques to estimate the sampling variability of *ũ*^(*i*)^. Recall that estimates of a network metric, *ũ*^(*i*)^, are a function of estimated networks 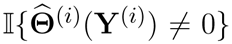. Unfortunately, closed form finite sample distributions for sparse penalized estimates of 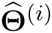 [**Berk et al.**, 2013] as well as sampling distributions of network metrics [**Balachandran et al.**, 2013] are still an emerging area of research. Thus, our problem differs from standard univariate GLM analyses employed in both voxel-wise activation studies and seed-based correlational analysis [**Penny et al.**, 2003; **Fox et al.**, 2006] where closed form asymptotic formulas for sample variance at the subject level are incorporated into the group level analyses. To tackle the issue of unknown sampling variability we build an empirical distribution of network statistics, where we perturb the data by sampling *m* out of *t* observations with replacement (bootstrap) [**Efron and Tibshirani**, 1993] or without replacement (subsampling) [**Politis et al.**, 1999] and re-estimate the network metrics per resample. By aggregating network statistics across resamples within each subject [**Breiman**, 1996a], we gain the additional benefit of variance reduction [Biihlmann and Yu, 2002] for individual subject metrics. Many variations of resampling techniques exist to handle dependencies [**Lahiri**, 2013] in spatio-temporal data. Since we assume approximately independent observations, from here on our resampling consists of sampling *t* out of *t* observations with replacement.

Recall that our method **R**^2^ is a variant of **R**^3^, that only involves resampling without random adaptive penalties. Here we obtain a bootstrapped network estimate 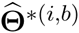, and a corresponding network metric *ũ*\*^(*i*,*b*)^ in Step 1 of our Algorithm 1 for each of *B* = 100 resamples. For our alternative procedure, **R**^3^, however, we not only use resampling, but simultaneously perturb the initial regularization parameters *λ*^(*i*)^ for every resample. This amounts to solving a weighted graphical lasso to re-estimate the network, where the weights are given by random adaptive penalties. Our motivation to use **R**^3^ is based on previous work in the context of two-sample tests for edge differences. **Narayan et al.**, [2015] show that random penalization significantly improved power over pure resampling to detect differential edges when the networks were moderately dense. Given this result, we sought to investigate the benefits of random penalization for more general network metrics. Intuitively, we anticipate that density based metrics beyond the edge level are immune to some graph selection errors. For instance, when false negatives are compensated by an equal number of false positive edges within the same node or subnetwork, node or subnetwork density values remain unchanged. However, graph selection errors that do not cancel each other out result in a net increase or decrease in density, thus contributing to loss of power. In these scenarios, we expect **R**^3^ to offer additional statistical power to test covariate effects.

Whereas general network metrics, require global properties of the network structure be preserved, the standard randomized graphical lasso [**Meinshausen and Buhlmann**, 2010] penalizes every edge randomly such that topological properties of the network could be easily destroyed within each resample. Thus, we seek to randomly perturb selected models in a manner less destructive to network structure. To achieve this, we adaptively penalize [**Zhou et al.**, 2011] entries of Θ^(*i*)^. Strongly present edges are more likely to be true edges and should thus be penalized less, whereas weak edges are more likely to be false and should be penalized more. As long as we have a good initial estimate of where the true edges in the network are, we can improve network estimates by adaptively re-estimating the network, while simultaneously using random penalties to account for potential biases in the initial estimates. In order to obtain a reliable initial estimate of network structure, we take advantage of the notion of stability as a measure of confidence popularized by **Breiman** [1996b]; **Meinshausen and Buhlmann** [2010]. Here the stability of an edge within a network across many resamples measures how strongly an is edge present in the network. When an edge belongs to the true network with high stability we randomly decrease the associated penalty by a constant *κ*. Conversely, we randomly increase the penalty by *κ* for an edge with low stability. Similar to **Narayan et al.**, [2015], we fix the constant *κ* to 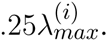. Here 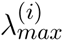 is the regularization parameter that results in the all zero graph for a subject. We call this approach random adaptive penalization (RAP) as it builds on the previous random penalization approach of **Narayan et al.**, [2015] but adaptively perturbs the regularization parameters using initial stability scores along the lines of the random lasso [**Wang et al.**, 2011].

Since, random adaptive penalization depends on an initial estimate of the stability of every edge in the network, we take advantage of the basic resampling step in Algorithm 1 to obtain a stability score matrix **п**̂^(*i*)^ for each subject. The entries of this matrix provide a proportion that takes values in the interval (0,1). Once we have the stability scores, we consider an additional set of *B* = 100 resamples to implement RAp. Thus in step 2 of Algorithm 1, we form an matrix of random penalties 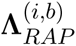 per resample *b*. For each edge (*k*, *l*) the corresponding adaptive penalty is determined by perturbing initial *λ̂*^(*i*)^ by an amount *κ* using a Bernoulli random variable. The probability of success of each Bernoulli r.v is determined by the corresponding stability score for that edge.

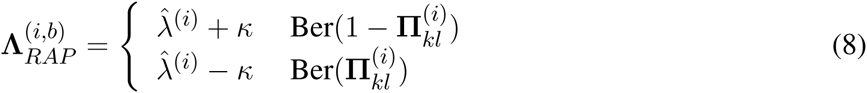

Putting these components together, **R**^3^ consists of first running Step 1 of Algorithm 1 to obtain stability scores and then using an additional *B* resamples based on random adaptive penalization, summarized in Step 2 of Algorithm 1 to obtain *nB* resampled network metrics *ũ*^(*i*,*b*)^. Note that in subsequent steps we omit the superscripts in 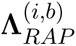 for notational convenience.

#### 2.3.3 Test Statistics for Network Metrics

Both **R**^2^ and **R**^3^ yield a total of *nB* resampled network statistics that possess two levels of variability. If we applied single level regression techniques to test the covariate effect in (3), we would in effect assume that all the *nB* resampled statistics were independent. Test statistics that assume *nB* independent observations, despite the availability of only *n* independent clusters of size *B* are known to be overoptimistic [**Laird and Ware**, 1982; **Liang and Zeger**, 1993]. To address this overoptimism, a more reasonable assumption is that resampled statistics between any two subjects are independent, whereas within subject resampling statistics are positively correlated. Just as we commonly employ mixed effects models to account for two levels of variation in repeated measures data, we employ similar two-level models to derive test statistics for resampled network metrics.

Let 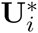 denote the vector *B* × 1 vector of resampled statistics per subject {*ũ*\*^(*i*,*b*)^} In the case of real valued density metrics, we use a linear mixed effects (LME) model for repeated measures [**Laird and Ware, 1982**] to account for the two levels of variability in resampled statistics.

#### Algorithm 1 : **R**^2^ and **R**^3^ Procedures for Testing Covariates Effects on Network Metrics

*Step 0:* **Initial Parameters**

**Input: Y**^(*i*)^, **Output:** *λ̂*^(*i*)^

Estimate *λ̂*^(*i*)^ using graphical model estimation and selection (StARS) for each subject *i*.

*Step 1:* **Subject Level Resampling**

**Input:** (**Y**^(*i*)^, *λ̂*^(*i*)^, **B** = 100), **Output:** Either *ũ*\*^(*i*,*b*)^ or **п**̂^(*i*)^

(a) FOR *b* = 1,…, *B* in the *i*^*th*^ subject

(i) Bootstrap the data **Y**^(*i*)^ to get **Y**\*^(*i*,*b*)^ and sample correlation matrix *Σ̃*\*^(*i*,*b*)^
(ii) Perform a standard graphical lasso 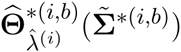 in Eq. (7)
(iii) **If R**^2^: Compute network statistic *ũ*\*^(*i*,*b*)^ defined in Section 2.1 END
(b) **If R**^3^: Estimate stability scores 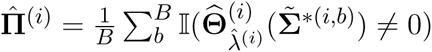

*Step 2:* **Subject Level Resampling & Random Adaptive Penalization (R**^3^ **only)**

**Input:** (**Y**^(*i*)^, **п**̂^(*i*)^, *λ̂*^(*i*)^,*B* = 100), **Output:** *ũ*\*^(*i*,*b*)^

(a) FOR *b* = 1,…, *B* in the *i*^*th*^ subject
  (i) Bootstrap the data **Y**^(*i*)^ to get **Y**\*^(*i*,*b*)^ and sample correlation matrix Σ̃*^(*i*,*b*)^
  (ii) Using stability scores from Step 1(b), compute random adaptive penalties 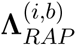 in Eq. (8)
  (iii) Using a weighted graphical lasso, estimate 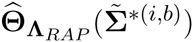 in Eq. (7)
  (iv) Compute network statistic *ũ*\*^(*i*,*b*)^ defined in Section 2.1

END

*Step 3:* **Population Level Inference for *β̂* using Random Effects Input:** 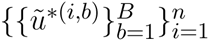, **Output:** *β̂* and *p*-values

(a) Estimate fixed covariate effects *β̂* using mixed effects models. (Section B.1)
(b) Compute mixed effects test statistic and *p*-values in Eq. (B.1)

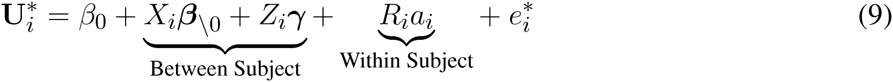

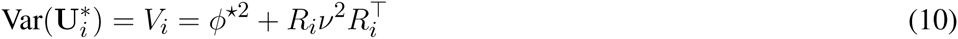

Here *a*_*i*_ are *i.i.d* subject level random intercepts with variance *Var*(*a*_*i*_) = *ν*^2^, *R*_*i*_ = 1_*B* × *i*_ is the random effect design matrix, and 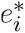 is independent of *a*_*i*_ and captures within subject sampling variability with variance *Var*(*e*_*i*_) = *ϕ*^⋆2^Ι_*B*_ where Ι denotes the identity. From hereon, we ignore the intercept *β*_0_, and assume that *β* denotes the (*q* × 1) vector of explanatory fixed effects.

Estimation and inference for linear mixed effect models are well covered in the neuroimaging literature in the context of functional activation studies and longitudinal designs [**Beckmann et al.**, 2003; **Bernal-Rusiel et al.**, 2013]. We employ standard estimators and test statistics for linear mixed effects models including generalized least squares estimators for *β̂* and corresponding restricted maximum likelihood (ReML) estimators of variance to obtain F-test statistics to test the null hypothesis regarding *β*,the covariate effects. A thorough review of mixed effects models can be found in **Agresti** [2015] and we also spell these out in more detail for our methods in supplementary materials.

## 3 SIMULATION STUDY

IIn this section, we seek to evaluate our framework for testing covariate effects by conducting a rigorous power analysis using realistic fMRI network structures. We obtain realistic network structures for fMRI functional connectivity by using networks estimated from real data as the basis of our simulated networks. First, we synthetically create multivariate data according to our two-level models using realistic graph structures in Section 3.1. Since we know the true structure of graphical models and their network metrics we empirically measure statistical power and type-I error for all methods. Then, in Section 3.2 we offer two key results. First, by employing simulations using two-level models of variability in (4) that reflect how functional networks are analyzed in practice, we provide a more realistic assessment of when we lose statistical power due to sample sizes (*t*, *n*) and covariate signal-to-noise (SNR) controlled by population variance *ν*^2^. Second, we show that both **R**^2^ and **R**^3^ mitigate the challenges discussed in Section 2.2 and improve statistical power over standard test statistics under various sample sizes and covariate SNR regimes.

### 3.1 SIMULATION SETUP FOR NODE AND SUBNETWORK DENSITY

We simulate multivariate data according to our two level models in Section 2.1. We know from previous work that the graph structure or location of non-zeros in the inverse covariance [**Narayan et al.**, 2015] influences the difficulty of estimating individual subject networks accurately. Using a group level empirical inverse correlation matrix obtained from 90 healthy subjects in the Michigan sample of the ABIDE dataset, preprocessed in Section 4, we threshold entries smaller than *τ* = |.25| to create a baseline network A_0_ that contributes to the intercept term *β*_0_ of our model (4). Illustrations of this baseline network can be found in Figure A.0 in the supplementary materials. Then we create individual adjacency matrices and network metrics *u*^(*i*)^ according to the linear model (5). We create inverse correlation matrices Θ^(*i*)^ using the graph structure provided by *A*_0_ and ensure Θ^(*i*)^ is positive definite.

Our main focus in the simulation study is to conduct a rigorous power analysis to detect covariate effects on node density and subnetwork density under a range of sample sizes and population variability and demonstrate the benefits of using **R**^3^ and **R**^2^ over standard approaches. Recall from Section 2.1 that node density is the degree of a node, while the subnetwork density is the number of connections between setsof nodes that make up a submatrix or subnetwork of the inverse covariance matrix. We obtain empirical estimates of statistical power by measuring the proportion of times we successfully reject *β̂*_\0_ = 0 at level *α* = .05, in the presence of a true covariate effect *β*_\0_ = 0, across 150 monte-carlo trials for a simulation scenario. Similarly, we obtain an empirical estimate of type I error by measuring the proportion of times we reject *β̂*_\0_ = 0 at level *α* = .05 in the presence of a null covariate effect of *β*_\0_ = 0.

Although one could choose to vary a large number of parameters for these simulations, we focus on the parameters most important for a power analysis, sample sizes and population variance, (*t*, *n*, *ν*^2^), while fixing other parameters such as number of covariates to *q* = 1, *r* = 0 and number of nodes to *p* = 50. We present a 3 × 3 panel of 9 power analyses of node density in Figure 2 where we vary *t* = {*p*, 2*p*, 4*p*} along the y-axis and *ν*^2^ = {.1, .25, .5} along the x-axis. Then within each sub-panel, we evaluate statistical power at subject sample sizes of *n* = {5,10,… 95}. For the entire 3 × 3 panel we hold the intercept and covariate effect fixed at *β*_0_ = 2, *β*_1_ = 1. Thus, each sub-panel illustrates statistical power as a function of subject sample size *n* for a fixed value of (*t*, *ν*^2^). Similarly, in Figure 3 we present power analyses for subnetwork density where we hold the intercept and covariate fixed at *β*_0_ = 5, *β*_1_ = 2 and use subnetworks of size .1*p* = 10 nodes. We use larger values for covariate effects to ensure that the number of edges in a subnetwork are realistically large for a subnetwork with 10 nodes. While the sample sizes (*t*, n) are identical to those in node density, we increase *ν*^2^ = {.4,1, 2} to match *β*. This ensures that covariate signal to noise ratio 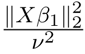 is similar for both metrics. Note that that the intercept values *β*_0_ in both power analyses were based on the average node degree in *A*_0_ or average subnetwork density for subnetworks of size 10 in *A*_0_. For each power analysis, we have a corresponding simulation of type-I error, obtained by setting *β*_1_ = 0 while keeping all other parameters equivalent. The full set of type-I error control results are presented in supplementary materials, and one representative simulation for each metric is presented in Figure 4.

**Figure 2.**
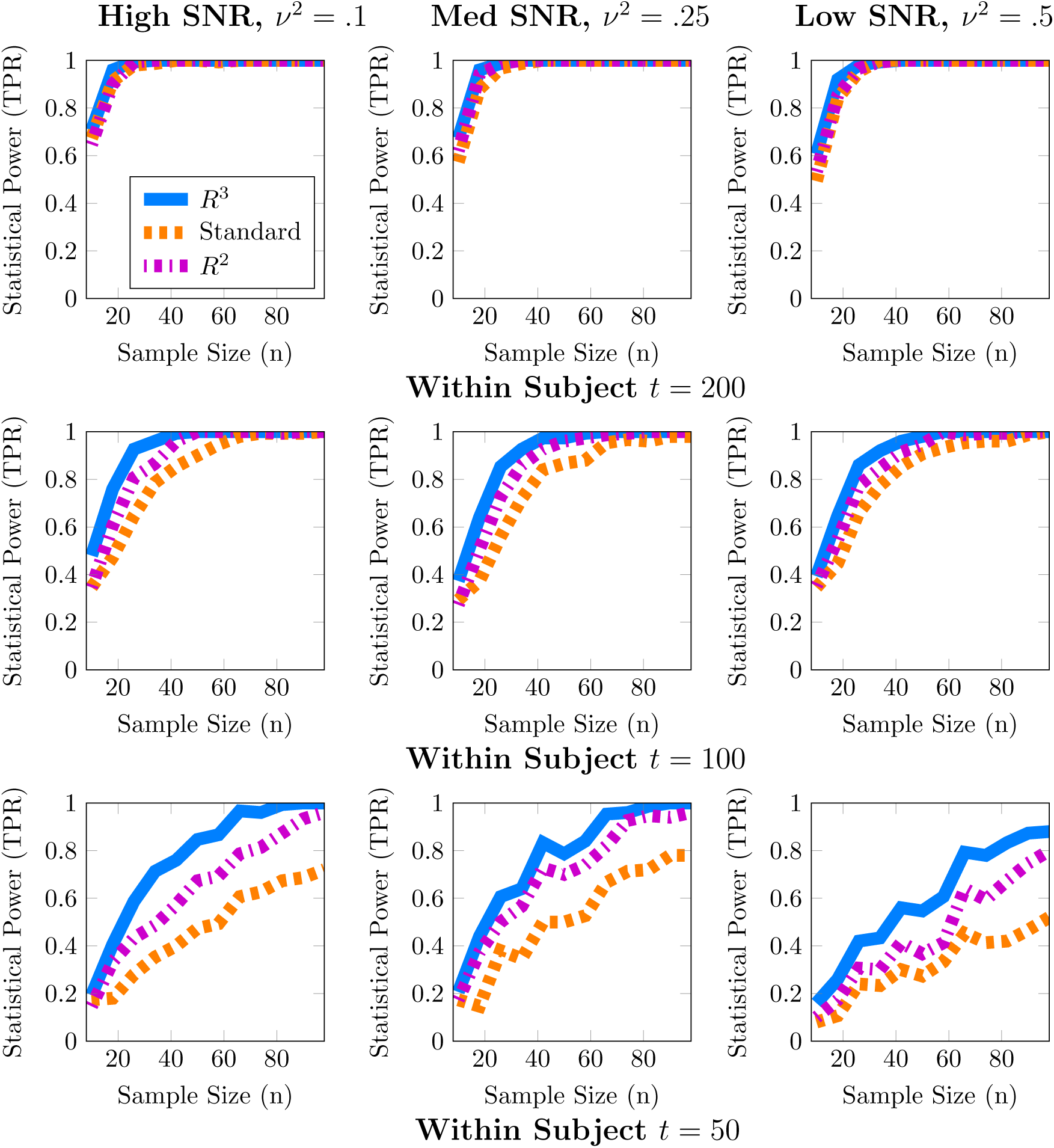
Statistical Power Analysis for Node Density. When node density varies with an explanatory covariate (*q* = 1), statistical power to detect this covariate effect improves with subject sample size *n* but crucially depends on the number of independent fMRI samples *t* from a single session and relative size of the covariate effect, *β*_1_ = 1, to population variance *ν*^2^ (covariate SNR). When *t* ≈ *p*, estimates of node density are both highly variable and potentially biased. By accounting for these issues, **R**^3^ and **R**^2^ improve estimates of network metrics, thus exceeding 80% power, whereas the standard F-test is substantially less powerful. Note that **R**^3^ and **R**^2^ are more powerful at smaller sample sizes compared to the standard approach. However, when fMRI samples become sufficiently large at *t* ≈ 4*p*, all methods become similarly powerful for detecting covariate effects of node density. Empirical statistical power is 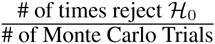 when the alternative is true in (3).

**Figure 3.**
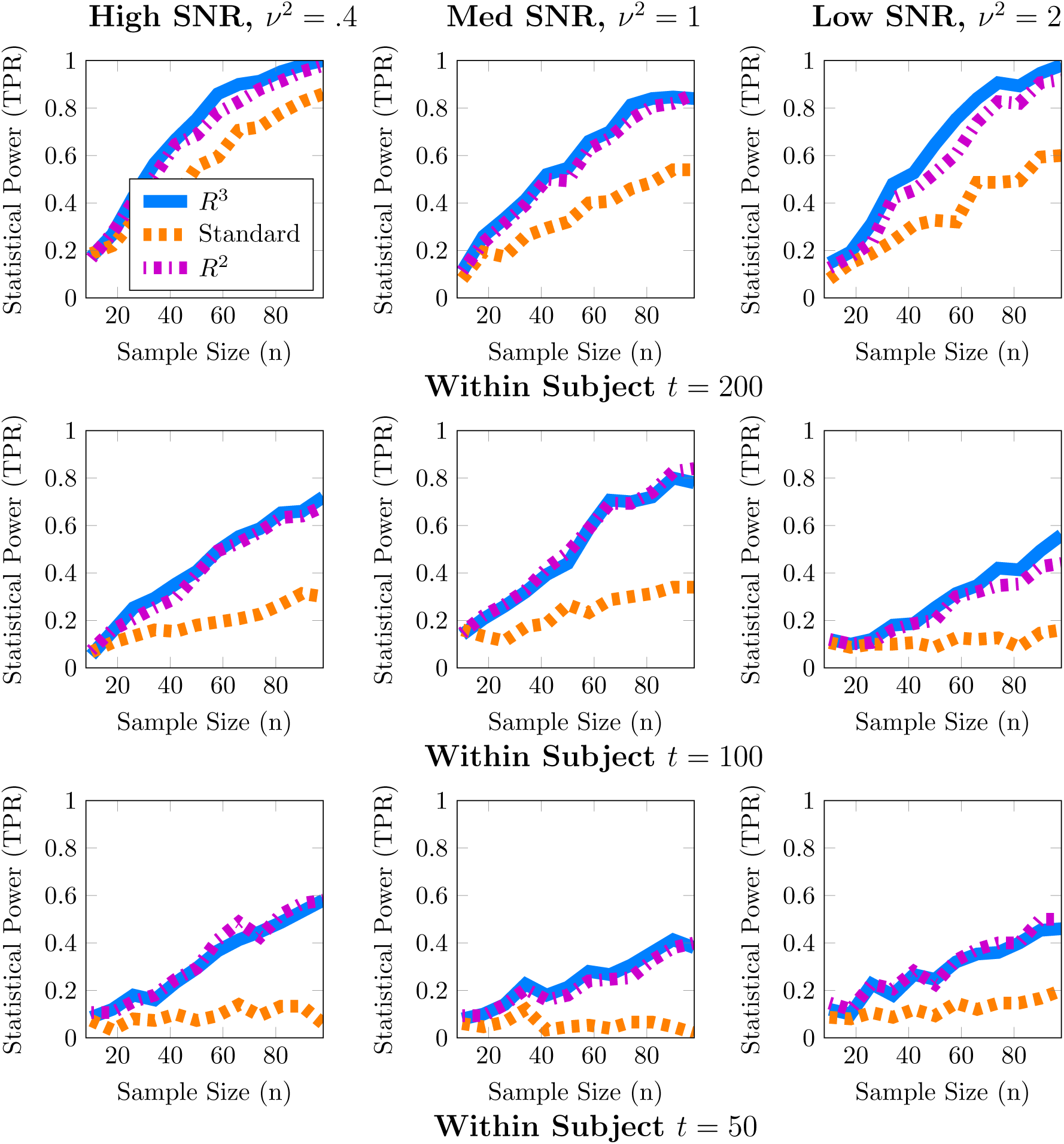
Statistical Power for Subnetwork Density. When subnetwork density varies with an explanatory covariate (*q* = 1), statistical power to detect this effect improves with subject sample size *n* but crucially depends on the number of independent fMRI samples *t* from a single session and the relative size of the covariate effect, *β*_1_ = 2, to the population variance *ν*^2^ (covariate SNR). For many values of (*t*,*p*) estimates of subnetwork density are both highly variable and potentially biased. By accounting for these issues, both **R**^3^ and **R**^2^ test statistics substantially improve statistical power across all regimes at smaller subject sample sizes, whereas the standard F-test is substantially less powerful. We note that covariate effects on subnetwork metrics are particularly hard to detect when *t* ≈ *p*, with statistical power often below 60%. Empirical statistical power is defined as 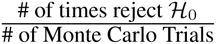 when the alternative is true in (3).

**Figure 4.**
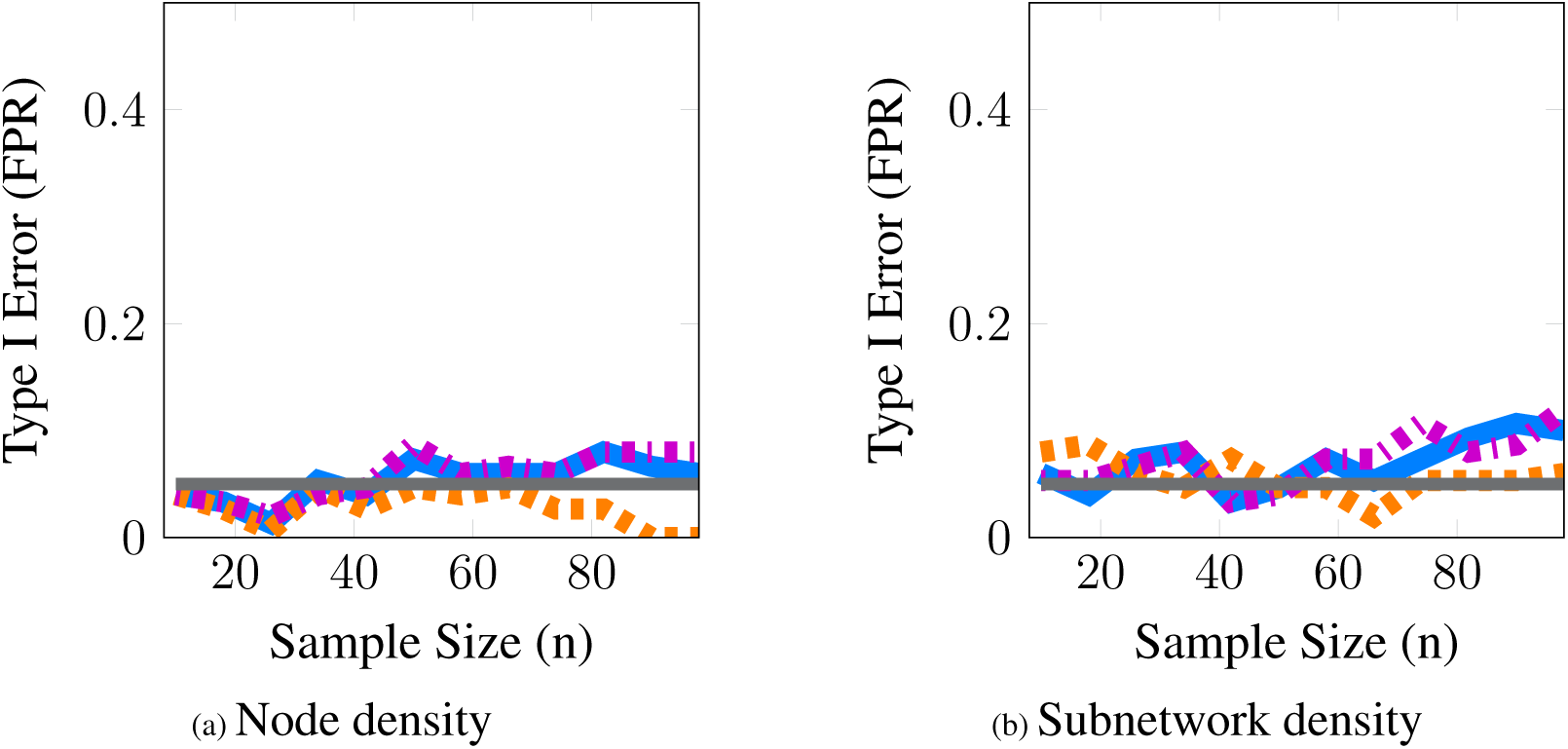
Statistical Type I Error Is Controlled for both Node and Subnetwork Density. These simulations evaluate the level of our tests; we report the estimated type-I error as a function of subject sample size *n*. The grey line represents the 5% level of the test. Here, we provide a representative simulation for node and subnetwork density in the moderate SNR regime with (*p* = 50,*t* = 100) and *ν*^2^ = .25 for node density and *ν*^2^ = 1 for subnetwork density. All methods approximately control type Ι error across all scenarios studied for both metrics. The full panel of simulations that complement the power analyses in Fig 2 and Fig 3 are included in supplementary materials.

### 3.2 SIMULATION RESULTS

In these simulations, our methods, **R**^3^ and **R**^2^, empirically outperform standard methods in terms of statistical power, particularly when within subject observations are comparable to the dimension of the network, and subject networks are harder to estimate correctly. Recall from Section 2.2 that we expect to lose statistical power when individual subject networks are difficult to estimate correctly, due to additional sampling variability and bias in network metrics. As expected, power analyses for both metrics in Figures 2 and 3 reveal that statistical power deteriorates as observations *t* available for subject network estimation reduces. Moreover, this loss of statistical power cannot always be compensated by larger subject sample sizes *n*. For example, the best achievable statistical power at large subject samples of *n* ≈ 100 begins to deteriorate when *t* = *p*. While, the best achievable statistical power often exceeds 90% for node density when *t* > *p*, it drops as low as 80% for **R**^3^ and **R**^2^. The standard approach in contrast drops below 60% node density. In the case of subnetwork density, statistical power for **R**^3^ and **R**^2^ exceed 80% when *t* = 4*p*, this drops as low as 60% at more modest sample sizes of *t* = 2*p* and further down to 40% at *t* = *p*. The standard approach falls to below 40% more quickly at *t* = 2*p* and below 20% when *t* = *p*.

Just as with subject sample size, when individual network estimation is easy in our simulations with larger within subject observations of *t* = 4*p*, the covariate signal to noise ratio or SNR has an almost negligible impact on statistical power. However, as *t* decreases, network estimation becomes harder and consequently, all methods become much more sensitive to SNR. For example, in regimes where *t* = 2*p*, network estimation is moderately hard but detecting covariate effects is achievable at high SNR. However, we observe that all methods lose power as covariate SNR decreases. We also observe that loss of statistical power due to SNR is more pronounced at smaller subject sample sizes of *n* < 60. Such a result is expected since sampling variability of covariate effect *β*_1_ is proportional to population variance *ν*^2^ and decreases with larger subject sample sizes *n*.

We noted earlier in Section 2.3 that we expect the benefits of **R**^3^ over **R**^2^ to be the greatest for finest scale metrics at the edge level which are most sensitive to graph selection errors and decrease as metrics measure density at more global levels. Whereas random penalization improves statistical power relative to **R**^2^ for two-sample differences at the edge level **Narayan et al.**, [2015], they share similar statistical power for node and subnetwork density in most simulations presented here, with some marginal benefits for node density. **R**^3^ offers greater benefits over **R**^2^ at small sample sizes *t* for networks that are more sparse and where the stability of true edges over false edges can be improved via random penalties. All methods, including **R**^3^ and **R**^2^ are unable to detect covariate effects when estimation of individual networks becomes unreliable under high density regimes. We provide additional simulations that vary the sparsity of baseline networks in Figure A.3 in the supplementary materials.

Finally, in Figure 4, we provide evidence that type-I error is controlled by all methods for both node and subnetwork density. The full panel of simulations that complement Figures 2 & 3 are included in supplementary materials.

From these simulations we conclude that resampling based approaches are more efficient, i.e. they have higher statistical power for both node and subnetwork density at smaller subject sample sizes *n*, particularly for smaller *t* and lower covariate SNR. Another insight from these simulations is that given a fixed budget of fMRI session time, it is preferable to increase the number of within session observations *t* per subject for fewer number of subjects *n* in order to maximize statistical power. For studies where each fMRI session consists of observations comparable to the size of networks (*t*,*p* ϵ [100, 200]), as well as for studies that cannot recruit a large number of subjects, our methods, **R**^3^ and **R**^2^, make better use of available data and improve statistical power compared to standard approaches to network analysis.

## 4 CASE STUDY

A number of recent studies on autism spectrum disorders (ASD) have found differences in functional connectivity that were correlated with symptom severity as measured by Autism Diagnostic Interview (ADI) or Autism Diagnostic Observation Schedule (ADOS). However, the majority of these studies that link symptom severity to functional connectivity derive networks using pairwise correlations [**Uddin et al.**, 2013b; **Supekar et al.**, 2013]. An important shortcoming of studying differences in pairwise correlation networks is that edges in a true correlational network might be present due to the effect of “common causes” elsewhere in the brain and do not necessarily represent a direct flow of information. Thus, while correlational networks can provide network biomarkers for autism [**Supekar et al.**, 2013], it is more problematic to infer network mechanisms of behavioral deficits in ASD exclusively using correlational networks. However, by studying previously implicated regions and subnetworks using Gaussian graphical models (GGMs), we strengthen the interpretation of variations in network structure linked to autism severity. Thus, by employing our two level models (1) based on GGMs to detect covariate effects, we enable scientists to infer that any network differences linked with behavioral deficits implicate nodes and edges directly involved in the disease mechanism. Guided by the successes of our simulation study, we employ **R**^3^ to investigate the relationship between cognitive scores on node and subnetwork densities in autism spectrum disorders. In particular, we conduct tests for covariate effects on two density metrics, the node density and subnetwork density. Node density counts the number of connections between a single region of interest to all other regions where as subnetwork density counts the number of connections between sets of regions or subnetworks. We investigate nodes and subnetworks hypothesized in the literature [**Uddin**, 2014] to be involved in regulating attention to salient events and explanatory for behavioral deficits in ASD.

### 4.1 ABIDE DATA COLLECTION AND PREPROCESSING

We use resting state fMRI data collected from the Autism Brain Imaging Data Exchange (ABIDE) project [**Di Martino et al., 2014b**] and preprocessed by the Preprocessed Connectomes Project (PCP) [**Craddock and Bellec**, 2015] using the configurable-pipeline for analysis of connectomes or (C-PAC) toolbox [**Craddock**, 2014; **Giavasis**, 2015]. In order to properly account for site effects, we choose to focus on two major sites with relatively large samples, UCLA and Michigan, resulting in 98 and 140 subjects per site. While both ADOS and ADI-R cognitive scores are available for these sites, we focus on ADOS scores obtained using the Gotham algorithm [**Gotham et al.**, 2009], which is known to be comparable across different age groups.

The ABIDE data was acquired [**Di Martino et al.**, 2014b] using T2 weighted functional MRI images with scan parameters TR= 2 at the Michigan site and TR= 3 at the UCLA site. Subsequently, this data was minimally preprocessed using the C-PAC utility [**Giavasis**, 2015; **Craddock and Bellec**, 2015], including slice timing correction, motion realignment and motion correction using 24 motion parameters, and normalization of images to Montreal Neurological Institute (MNI) 152 stereotactic space at 3 × 3 × 3 mm^3^ isotropic resolution. The pipeline was also configured to regress out nuisance signals from the fMRI time-series. The nuisance variables included were physiological confounds such as heart beat and respiration, tissue signals and low frequency drifts in the time-series. We did not regress out the global signal as this operation is known to introduce artifacts in the spatial covariance structure [**Murphy et al.**, 2009]. Additionally, we did not apply band pass filtering as this would interfere with subsequent temporal whitening that we describe later in thisSection. Preprocessed data without bandpass filtering and global signal regression is available using the *noglobalnofilt* option in the PCP project. Finally, the spatial time-series was parcellated into times-series × regions of interest using the Harvard-Oxford atlas distributed with FSL http://fsl.fmrib.ox.ac.uk/fsl/fslwiki/. Here we included *p* = 110 regions of interest including 96 cortical regions and 14 subcortical regions. Regions corresponding to white matter, brain stem and cerebellum were excluded. The resulting time-series × regions data matrix for each individual subject is (*t* = 116, *p* = 110) for UCLA subjects and (*t* = 300, *p* = 110) for Michigan subjects. This preprocessed dataset has been archived in a public repository http://dx.doi.org/10.6084/m9.figshare.1533313.

### 4.2 PREVIOUSLY IMPLICATED SUBNETWORKS AND REGIONS

Distinct lines of evidence suggest the involvement of limbic, fronto-parietal, default mode and ventral attention regions in ASD. Uddin [2014] summarize the evidence in favor of a salience-network model to explain behavioral dysfunction in responding to external stimuli. According to this model, the salience network regions that span traditional limbic and ventral attention systems play a vital role in coordinating information between the default mode regions involved in attending to internal stimuli and the frontoparietal regions involved in regulating attention to external stimuli. Together, these interactions enable appropriate behavioral responses to “salient” or important events in the external environment. **Uddin et al.**, [2013a] conducted a network-based prediction study and found that connectivity features of the anterior cingulate cortex, and the anterior insula, predict an increase ADOS repetitive behavior scores. Similarly, another study by **DiMartino et al.**, [2009] also implicates connectivity of anterior insula and anterior cingulate cortex to deficits in social responsiveness in autism. **Cherkassky et al.**, [2006]; **Monk et al.**, [2009] implicate posterior cingulate connectivity within the default mode network in ASD. **Alaerts et al.**, [2014] show that deficits in emotion recognition were correlated with network features in the right posterior superior temporal sulcus, a result also supported in the wider literature [**Uddin et al.**, 2013b].

Additionally, we also major findings from previous analyses of the ABIDE dataset that include the UCLA or Michigan subject samples. Whole brain voxelwise analysis by **Di Martino et al.** [2014b] revealed covariate effects associated with the mid insula, posterior insula, posterior cingulate cortex and thalamus. Group level two-sample tests of functional segregation and integration in seed based functional connectivity [**Rudie et al.**, 2013, **2012**] reveal differences in the amygdyla, IFG right pars opercularis.

Based on our review of existing literature, we seek to detect covariate effects with respect to 23 hypotheses regarding the density of connections. Of these 23 hypotheses, 13 correspond to density of connections of nodes or brain regions with respect to the whole brain, and 10 correspond to the density within and between 4 large scale functional subnetworks. These regions are defined using the Harvard-Oxford atlas with large scale subnetworks provided by **Yeo et al.**, [2011]. Figure 6 illustrates the volumes associated with the 13 regions of interest. Figure 5 illustrates the four large scale functional brain networks we consider, namely, the default mode, the frontoparietal, the limbic and the ventral attention networks as defined by **Yeo et al.**, [2011]. By explicitly testing the density of long-range connections in brain regions and networks previously linked with ASD, we aim to identify network structures at the node and subnetwork level that are directly involved in behavioral deficits.

**Figure 5.**
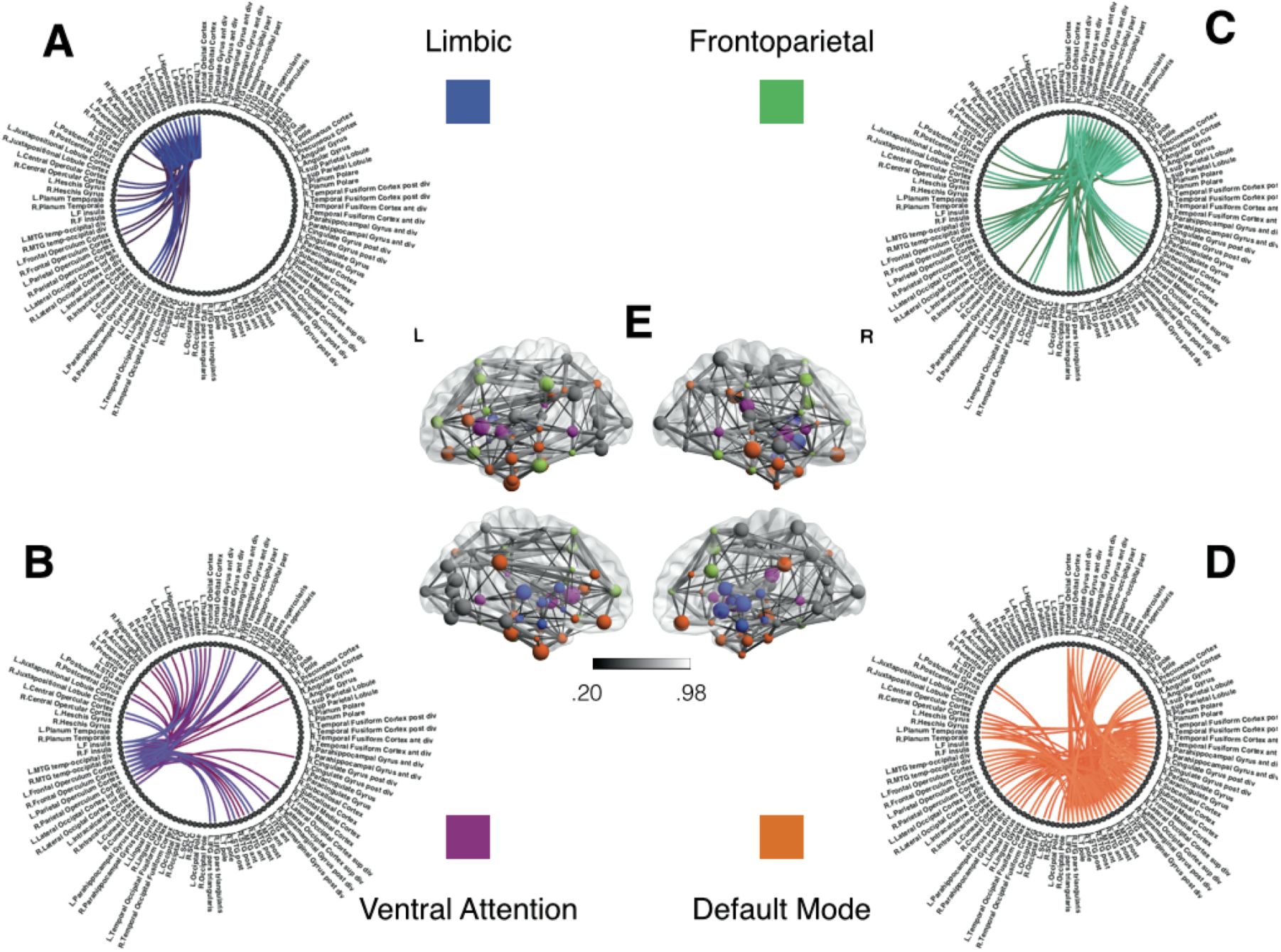
Functional Subnetworks of Interest for Covariate Tests of Network Density. This figure illustrates the subnetworks we have chosen to test for covariate effects in Table 1. Using previous studies discussed in 4.2, we seek to test whether symptom severity is associated with individual differences in the density or number of connections within and between these sub-networks. Panels A-D illustrate subnetwork components of the full group level network in panel E. The network structure in Panel (A) shows links within the limbic subnetwork as well as between the limbic regions and all other brain regions. Similarly, each of the other panels emphasize connectivity of fronto-parietal (B), ventral attention (C) and default mode (D) regions, respectively, to the whole brain. For the purposes of illustration, this group level network is obtained using individually estimated graphical models from the procedure in Section 2.3.1. Nodes correspond to anatomical regions in the Harvard Oxford Atlas [**Fischl et al.**, 2004]. The subnetworks correspond to resting state networks provided by **Yeo et al.**, [2011]. We first threshold weak edges with stability scores less than .8 in individual subject networks and then obtain a group level network by aggregating edge presence across all subjects. Note that we use this group network exclusively for illustrative purposes and not for statistical inference. The color gradient for edges in group network in panel E corresponds to proportion of stable edges found across all subjects.

**Figure 6.**
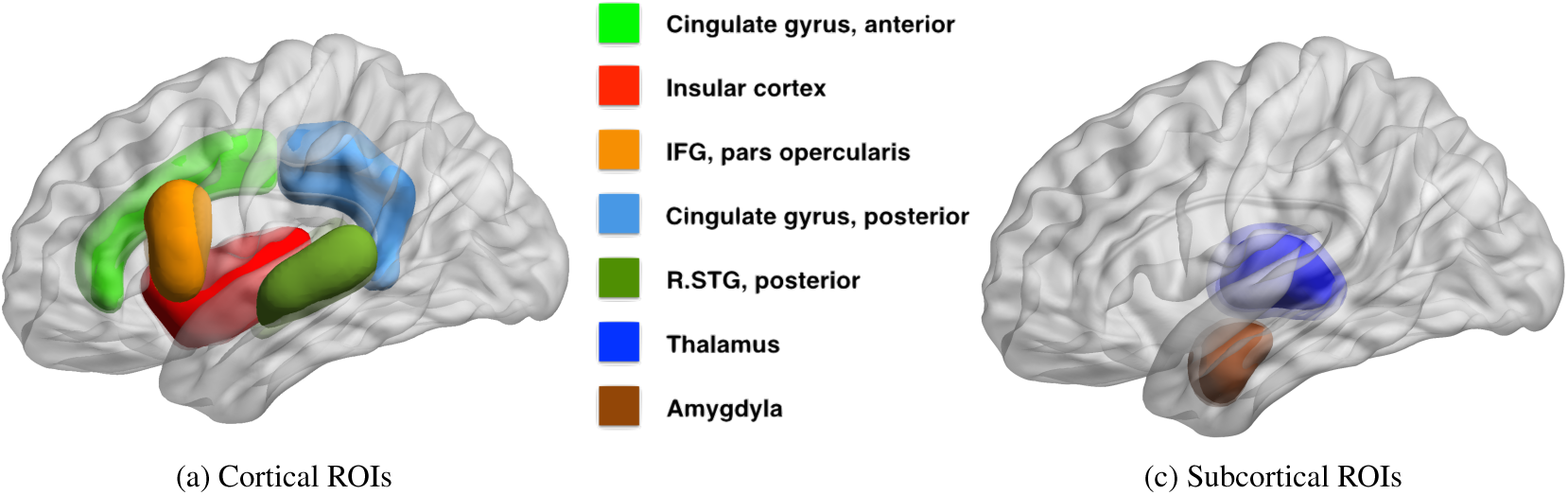
Regions of Interest for Covariate Tests of Node. Density. This figure illustrates the regions of interest based on the Harvard Oxford Atlas that we have chosen to test for covariate effects in Table 2. Several studies link the severity of autism spectrum disorders, measured by ADI or ADOS cognitive scores, with 9 cortical (Fig 6a) and 4 sub-cortical (Fig 6c) regions of interest, all within the default mode, limbic, frontoparietal and ventral attention networks. The full literature review is available in 4.2.

#### 4.2.1 Testing for covariate effects via **R**^3^

We employ the linear model from (5) for node and subnetwork density to test the null hypothesis that ADOS covariates have no effect on density. For this analysis, we jointly consider two related explanatory covariates, the ADOS Social Affect (SA) and the ADOS Restricted, Repetitive Behavior (RRB) scores (*q* = 2), while accounting for differences in clinical evaluation across sites, by incorporating site as a nuisance covariate (*r* = 1). We eliminate subjects without ADOS cognitive scores, leaving us with *n* = 100 autism subjects. Thus, the final data tensor for covariate tests contains either *t* = 116 (UCLA) or *t* = 300 (Michigan) time-points for *p* =110 brain regions in *n* = 100 subjects.

Before applying the **R**^3^ procedure from Section 2.3 to the preprocessed ABIDE dataset, we need to ensure fMRI observations are approximately independent. By whitening temporal observations, we ensure that estimating individual subject networks is more efficient. We achieve this by first estimating the temporal precision matrix 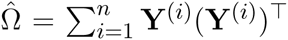 using the banded regularization procedure of [**Bickel and Levina**, 2008] for autoregressive data and whitening the fMRI time-series of each subject **Ỹ**^(*i*)^ = Ω̂ ^1/2^**Y**^*i*^. To choose the number of lags, we conduct model selection via cross-validation [**Bickel and Levina**, 2008]. Given these whitened observations, we apply the **R**^3^ procedure outlined in Algorithm 1. We initialize regularization parameters using StARS and subsequently perturb these parameters according to RAP as described in Section 2.3. Since we have a total of 23 node density and subnetwork density hypotheses, we control the false discovery rate at the 5% level using the Benjamini-Yekutieli procedure [**Benjamini and Yekutieli**, 2001].

### 4.3 ABIDE DATA ANALYSIS: RESULTS

Tables 1 & 2 show statistically significant covariate effects for 3 subnetwork hypotheses and 5 regions of interest. Notable findings amongst subnetwork hypotheses in Table 1 are that an increase in behavioral deficits indicated by restricted and repetitive behavior scores (RRB) and social affect (SA) is associated with a decrease in connection densities in frontoparietal-based subnetworks. The 3 prominent findings involve connection densities between the frontoparietal to limbic subnetworks, between the frontoparietal to ventral attention subnetworks and between the default mode and limbic subnetworks. Individual regression coefficients and confidence intervals for RRB and SA suggest that of the two covariates, RRB scores particularly dominate the decrease in subnetwork density for two of these results, particularly the frontoparietal-limbic subnetwork. The most prominent results amongst region of interest hypotheses in Table 2 suggest that ADOS symptom severity is again associated with hypoconnectivity or a decrease in the number of connections between each of the following regions with the rest of the network — bilateral pairs of anterior cingulate cortex (ACC); left posterior cingulate *cortex*(PCC); the right inferior frontal gyrus (IFG); and the thalamus. Note that we use a conservative Benjamini-Yekutieli procedure [**Benjamini and Yekutieli**, 2001] to control for FDR at the 5% level under arbitrary dependence amongst the 23 hypotheses tested. Under a less conservative procedure, Benjamini-Hochberg [**Benjamini and Hochberg**, 1995], four additional hypotheses including the within-frontoparietal subnetwork and the right PCC are statistically significant at 5% FDR control. While the regression coefficients for site effects are non-zero in both analyses, most confidence intervals either contain zero or are very close to zero and not statistically significant. The one exception amongst our prominent findings, the right ACC, shows statistically significant site effects. We also find site effects for two hypotheses where we did not detect ADOS effects, namely, the limbic to ventral attention subnetwork and right insula. However, these site effects are not statistically significant after correcting for multiplicity.

**Table 1.** Joint ADOS Covariate Effects on Subnetwork Density. We jointly test the effects of two ADOS covariates on subnetwork density while accounting for site effects as a nuisance covariate. Here, the most prominent findings suggest that a decrease in the number of direct connections between frontoparietal to limbic, between frontoparietal to ventral attention subnetworks and between default to limbic subnetworks is linked with increased ADOS symptom severity. This result is consistent with the hypothesis that abnormalities within the salience network, comprising anterior cingulate cortex (a region within our frontoparietal network) and insula (a region within our ventral attention network), results in a failure to regulate between attention to external stimuli versus attention to internal thoughts. A total of three subnetworks, denoted by *, survive corrections for multiplicity, using false discovery control over all 23 hypotheses tested at the 5% level using Benjamini-Yekutieli. Although estimates of site effects were non-zero, individual confidence intervals for most site-effects were close to zero and were thus not statistically significant after corrections for multiplicity. Results are discussed further in Section 4.3

| Subnetwork 1 | SubNetwork 2 | pval (RRB + SA) | RRB | CI (L) | CI (U) | SA | CI (L) | CI (U) | SITE | CI (L) | CI (U) |
| --- | --- | --- | --- | --- | --- | --- | --- | --- | --- | --- | --- |
| Default | Default | 0.061200 | -2.66 | -7.10 | 1.78 | -0.66 | -2.92 | 1.59 | 0.22 | -5.85 | 6.30 |
| Default | Frontoparietal | 0.010000 | -2.34 | -4.97 | 0.29 | -0.18 | -1.52 | 1.15 | -0.52 | -4.12 | 3.08 |
| Default | Limbic | 0.004530* | -1.51 | -3.06 | 0.04 | -0.10 | -0.89 | 0.69 | -0.30 | -2.42 | 1.82 |
| Default | Ventral Attention | 0.038000 | -0.83 | -1.76 | 0.10 | 0.08 | -0.40 | 0.55 | 0.37 | -0.91 | 1.64 |
| Frontoparietal | Frontoparietal | 0.007030 | -1.36 | -3.23 | 0.52 | -0.51 | -1.47 | 0.44 | -0.47 | -3.04 | 2.10 |
| Frontoparietal | Limbic | 0.000088* | -1.15 | -1.98 | -0.31 | 0.00 | -0.43 | 0.43 | 0.23 | -0.92 | 1.38 |
| Frontoparietal | Ventral Attention | 0.003793* | -0.61 | -1.16 | -0.06 | 0.03 | -0.25 | 0.31 | 0.75 | 0.00 | 1.50 |
| Limbic | Limbic | 0.530000 | -0.19 | -1.70 | 1.32 | -0.35 | -1.11 | 0.42 | -0.77 | -2.83 | 1.29 |
| Limbic | Ventral Attention | 0.955000 | 0.01 | -0.45 | 0.46 | -0.05 | -0.28 | 0.18 | -0.69 | -1.31 | -0.06 |
| Ventral Attention | Ventral Attention | 0.196000 | -0.05 | -0.50 | 0.40 | -0.21 | -0.44 | 0.02 | -0.24 | -0.86 | 0.37 |

**Table 2.**
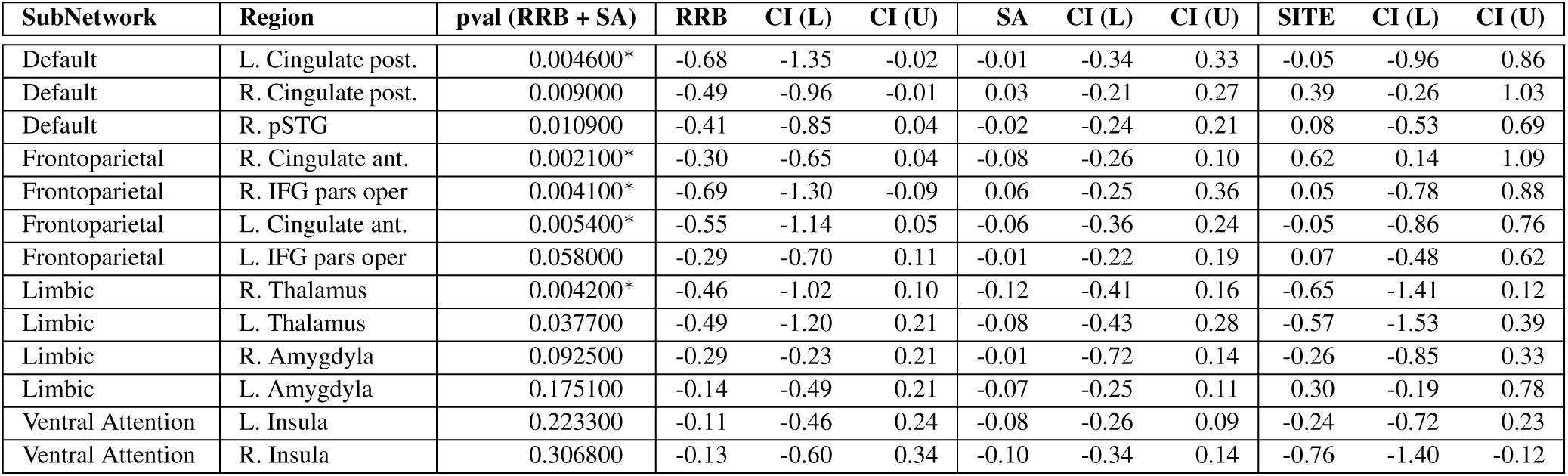
Joint ADOS Covariate Effects on Node Density. We jointly test the effects of two ADOS covariates on node density while accounting for site effects as a nuisance covariate. Notably, we find that a decrease in the number of direct connections between left posterior cingulate cortex (PCC) and anterior cingulate cortex (ACC) with all other regions is linked with an increase in ADOS symptom severity. This result corroborates previous findings that ACC (a component of the salience network) and PCC connectivity might be directly involved behavioral deficits ASD. A total of five regions, denoted by *, survive corrections for multiplicity, using false discovery control over all 23 hypotheses tested at the 5% level using Benjamini-Yekutieli. Although estimates of site effects were non-zero, individual confidence intervals for most site-effects were close to zero and were thus not statistically significant after corrections for multiplicity. Results are discussed further in Section 4.3

| SubNetwork | Region | pval (RRB + SA) | RRB | CI (L) | CI (U) | SA | CI (L) | CI (U) | SITE | CI (L) | CI (U) |
| --- | --- | --- | --- | --- | --- | --- | --- | --- | --- | --- | --- |
| Default | L. Cingulate post. | 0.004600* | -0.68 | -1.35 | -0.02 | -0.01 | -0.34 | 0.33 | -0.05 | -0.96 | 0.86 |
| Default | R. Cingulate post. | 0.009000 | -0.49 | -0.96 | -0.01 | 0.03 | -0.21 | 0.27 | 0.39 | -0.26 | 1.03 |
| Default | R. pSTG | 0.010900 | -0.41 | -0.85 | 0.04 | -0.02 | -0.24 | 0.21 | 0.08 | -0.53 | 0.69 |
| Frontoparietal | R. Cingulate ant. | 0.002100* | -0.30 | -0.65 | 0.04 | -0.08 | -0.26 | 0.10 | 0.62 | 0.14 | 1.09 |
| Frontoparietal | R. IFG pars oper | 0.004100* | -0.69 | -1.30 | -0.09 | 0.06 | -0.25 | 0.36 | 0.05 | -0.78 | 0.88 |
| Frontoparietal | L. Cingulate ant. | 0.005400* | -0.55 | -1.14 | 0.05 | -0.06 | -0.36 | 0.24 | -0.05 | -0.86 | 0.76 |
| Frontoparietal | L. IFG pars oper | 0.058000 | -0.29 | -0.70 | 0.11 | -0.01 | -0.22 | 0.19 | 0.07 | -0.48 | 0.62 |
| Limbic | R. Thalamus | 0.004200* | -0.46 | -1.02 | 0.10 | -0.12 | -0.41 | 0.16 | -0.65 | -1.41 | 0.12 |
| Limbic | L. Thalamus | 0.037700 | -0.49 | -1.20 | 0.21 | -0.08 | -0.43 | 0.28 | -0.57 | -1.53 | 0.39 |
| Limbic | R. Amygdyla | 0.092500 | -0.29 | -0.23 | 0.21 | -0.01 | -0.72 | 0.14 | -0.26 | -0.85 | 0.33 |
| Limbic | L. Amygdyla | 0.175100 | -0.14 | -0.49 | 0.21 | -0.07 | -0.25 | 0.11 | 0.30 | -0.19 | 0.78 |
| Ventral Attention | L. Insula | 0.223300 | -0.11 | -0.46 | 0.24 | -0.08 | -0.26 | 0.09 | -0.24 | -0.72 | 0.23 |
| Ventral Attention | R. Insula | 0.306800 | -0.13 | -0.60 | 0.34 | -0.10 | -0.34 | 0.14 | -0.76 | -1.40 | -0.12 |

Our analysis strongly implicates the frontoparietal-limbic subnetwork, and frontoparietal-ventral attention subnetworks, as well as posterior/anterior cingulate cortical connections with the rest of the brain, in behavioral deficits of ASD. Since we identify these regions and subnetworks using partial correlation measures of functional connectivity, our results provide strong evidence that these network components are directly involved in ASD. In particular, since the salience network [**Uddin et al.**, 2013a; **Buckner et al.**, 2013] is thought to comprise the ACC, which falls within our frontoparietal network, and insular regions that overlap limbic and ventral attention networks in our analysis, our subnetwork findings are consistent with the salience network explanation for behavioral deficits in autism. Additionally, our findings strongly implicate frontoparietal-limbic relationships. While our region of interest analysis found abnormalities in thalamar connectivity, a component of the limbic network, other limbic regions could also be directly involved in ASD and thus warrant further study.

We contrast our findings on the 23 a-priori hypotheses in Section 4.2 with previous analyses that were obtained by conducting network analyses on correlational networks, including previous analyses of the same ABIDE dataset. Our analysis detects only a subset of previous covariate effects on ASD network structure when using GGM based networks via **R**^3^. Correlational network analysis using the UCLA and UM samples of ABIDE [**Di Martino et al.**, 2014b; **Rudie et al.**, 2012] as well as those form alternative sites [**Uddin et al.**, 2013b] link insular, amygdylar connectivity with autism symptoms, whereas we do not detect strong effects for these regions for density metrics. The absence of strong covariate effects using GGMs suggests that the insular and amygdylar connections might be associated with behavioral deficits in autism only due to indirect correlations with other regions of interest. Similarly, although we find abnormalities in the PCC, a region within the default mode network, and between the default-mode and the limbic regions, we failed to find abnormalities linking the default mode with frontoparietal or ventral attention networks. This suggests that previous findings involving the default mode network could have been the result of indirect pairwise correlations, possibly driven by PCC and limbic regions. Although we use novel functional connectivity models and methods to analyze the ABIDE dataset, some of our choices of a-priori hypotheses for this analysis, notably, the inclusion of IFG pars opercularis and the amygdyla for node density, were guided by alternative analyses of the ABIDE dataset [**Rudie et al.**, 2012; **Di Martino et al.**, 2014b]. Thus, we need further validation of the purported effects of ADOS on IFG pars opercularis density.

## 5 DISCUSSION

This paper investigates an understudied issue in neuroimaging – the impact of (often imperfectly) estimated functional networks on subsequent population level inference to find differences across functional networks. Using an important class of network models for functional connectivity, Gaussian graphical models, we demonstrate that neglecting errors in estimated functional networks reduces statistical power to detect covariate effects for network metrics. While lack of statistical power due to small subject sizes is well documented in neuroimaging [**Button et al.**, 2013], recent test re-test studies [**Birn et al.**, 2013; **Laumann et al.**, 2015] suggest that typical fMRI studies of 5-10 minutes are highly susceptible to lack of statistical power. This paper provides additional evidence that within subject sample size, *t*, is important for well powered studies. For typical studies where *t* is comparable to the number of nodes *p*, errors in estimating functional networks can be substantial and not accounted for by standard test statistics. We show that our methods to mitigate this problem, **R**^2^ and **R**^3^, are always at least as powerful if not substantially more powerful than standard test statistics under a variety of sample sizes and covariate signal-to-noise regimes. Additionally, regardless of the methods employed, our power analyses suggest that in many scenarios, particularly when subject level networks are large, a more efficient use of a fixed experimental budget would be to collect more within subject measurements and fewer subject samples in order to maximize statistical power to detect covariate effects. While we demonstrate this result on the joint importance of within and between subject sample sizes using density based network metrics, we expect such results to hold more generally whenever population level functional connectivity analyses are conducted in a two-step manner where subject level networks are estimated initially and population level metrics then explicitly depend on the quality of subject level network estimates. In practice, we additionally need to incorporate other considerations beyond statistical power in choosing within subject scan length such as increase in movement or the discomfort to participants particularly in patient populations. These issues related to statistical power warrant further investigation in future work.

This paper also highlights the scientific merits of employing explicit density based metrics in graphical models of functional connectivity to gain insights into disease mechanisms at a macroscopic level using the ABIDE dataset [**Di Martino et al.**, 2014b]. In Section 4, we sought to detect covariate effects on the density of direct, long range functional connections in Austism Spectrum Disorders (ASD). Notably, our results in Section 4.3, at both the subnetwork and node level favor the hypoconnectivity hypothesis for behavioral deficits in ASD. Specifically, we find that a reduction in directly involved long-range functional connections between parcellated regions of interest increases ADOS symptom severity. Assuming that the salience network model of autism dysfunction is correct [**Uddin**, 2014], our results suggest that reduced interactions between the executive control network and the salience network, as well as default mode to the salience network might be responsible for ASD symptoms. Since we employ GGM based models, a plausible interpretation of such hypoconnectivity is that regions in ventral attention and limbic systems fail to adequately communicate with frontoparietal regions that participate in executive control and default mode regions that participate in internal attention. A previous study found evidence of hyperconnectivity when counting the number of local voxelwise connections in **Keown et al.**, [2013]. Our results do not contradict this finding since a network architecture of ASD could involve both reduced long range connections as well as increased density of local connections [**Rudie and Dapretto**, 2013]. Other results on hyperconnectivity [**Uddin et al.**, 2013a; **Supekar et al.**, 2013] do not explicitly employ degree or density of connections to measure hyper or hypo-conectivity but measure the strength of the mean pairwise correlation within and between regions and subnetworks. While the effect in **Supekar et al.**, [2013] appears to be a large and robust finding, the correlational model of connectivity employed in their analysis could be misleading since it includes both direct and indirect functional connections and does not explicitly measure the density of connections. While further studies are needed to resolve the questions raised by Rudie and Dapretto [2013] on this matter, we emphasize that since graphical models of functional connectivity capture direct functional connections, such models enable stronger scientific conclusions regarding functional network mechanisms compared to purely correlational models where edges do not necessarily reflect direct communication between regions.

As we discuss in the simulation results in Section 3.2, our ability to detect covariate effects in populations of graphical models deteriorates in highly dense regimes of network structure where the density or number of edges in the network increases substantially while the number of within subject observations remains limited, or when the individual networks contain a large number of hub-like structures[**Ravikumar et al.**, 2011; **Zhou et al.**, 2011]. Since our resampling based methods are a framework that employ existing graph estimation algorithms (Section 2.3), they inherit the strengths and limitations of the specific graph estimation algorithm in such high density regimes. By incorporating new and improved estimators [**Yang et al.**, 2014] for graphical models at the level of individual subjects, we expect corresponding variants of our resampling framework to detect covariate effects under a wider range of network density regimes.

While this paper specifically considers network models (1) where neuroimaging data is distributed according to a multivariate normal, alternative distributions can be employed for the subject level model in (1), including matrix variate distributions [**Allen and Tibshirani**, 2012; **Zhou**, 2014] that can account for the serial correlation in temporal observations, and nonparametric graphical models [**Lafferty et al.**, 2012] that relax assumptions of normality. Furthermore, while we focus on resting state functional connectivity in fMRI in this work, our concern regarding errors in estimating large functional networks is applicable to other imaging modalities including EEG/MEG studies. In fact, our two level models (1) and **R**^3^ framework can be easily extended to functional network analyses based on partial coherence [**Sato et al.**, 2009] networks or vector autoregressive models [**Koenig et al.**, 2005; **Schelter et al.**, 2006] that are popular in EEG/MEG studies. Additionally, our results are highly relevant to dynamic functional connectivity [**Chang and Glover**, 2010] analyses where studies estimate separate time-varying functional networks per subject using short sliding-windows of 30-60 seconds rather than 5-10 minutes. In such a high dimensional setting where *t* << *p*, our power analyses in Figures 2 and 3 suggest that such dynamic network analyses will be highly underpowered and could benefit from our methods. Thus, extensions of the **R**^3^ framework for dynamic connectivity analyses as well as other multivariate network models is a promising avenue of research. Other areas of investigation include inference for partial correlation strength and corresponding weighted network analysis, as well as including high dimensional covariates in our general linear model (2). Overall, this work reveals that accounting for imperfectly estimated functional networks dramatically improves statistical power to detect population level covariate effects, thus highlighting an important new direction for future research.

## 6 DATA AND SOFTWARE

The preprocessed ABIDE dataset used in this paper is available at http://dx.doi.org/10.6084/m9.figshare.1533313. Software for reproducing our analysis will be provided at https://bitbucket.org/gastats/monet.

## ACKNOWLEDGMENTS

The authors thank Steffie Tomson for helpful discussions and advice on preprocessing the ABIDE dataset.

*Funding:* M.N. and G.A. are supported by NSF DMS 1264058. M.N is supported an AWS (Amazon Web Services) research grant for computational resources.

## SUPPLEMENTARY MATERIALS

We include additional simulations and details of test statistics for our methods in the appendix.

### A SUPPLEMENTARY SIMULATIONS & FIGURES

In this appendix, we provide supplementary simulations and figures to complement the power analyses and summary of type-I error control that appear in Figures 3,4 & 5 of our manuscript. The setup for the supplementary simulations follows the procedures outlined in Section 4.1. Figures A.1 & A.2 provide a complete set of type-I error simulations for node and subnetwork density, respectively, and complement the power analyses found in Figures 3 & 4. Additionally, we demonstrate the impact of sparsity on the ability of **R**^3^, **R**^2^ and the standard method to detect covariate effects in Figure A.3. Here, we employ the node density metric in the medium SNR case (*ν*^2^ = .25) as a representative example, while holding all other parameters consistent with Figure 3 of the manuscript constant with exception of baseline sparsity threshold *τ*. While the simulations in our manuscript employed realistic networks (illustrated in Figure A.0) obtained by setting all partial correlations whose absolute values were less than *τ* = .25 to zero, we varied this threshold to values {.1, .4} to obtain both denser and sparser baseline networks.

**Figure A.0.**
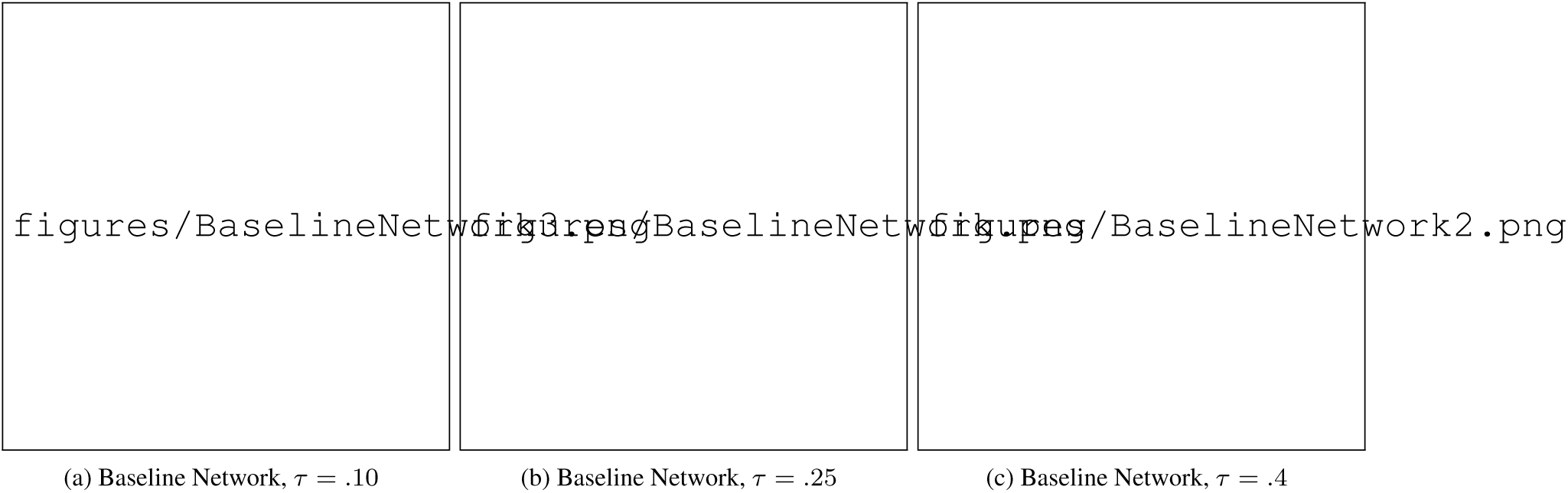
Empirically Derived Pseudo Real Networks. The simulations in the main paper in Figures 2 & 3 all begin with the baseline network (moderate density case) with threshold *τ* = .25 in (b). Then each individual subject network is simulated as described in Section 4.1. The simulations in Figures A.3 employ sparser and denser baseline networks given by thresholds *τ* = .1 and *τ* = .4 for the same experiment for node density in Figure 2.

**Figure A.1.**
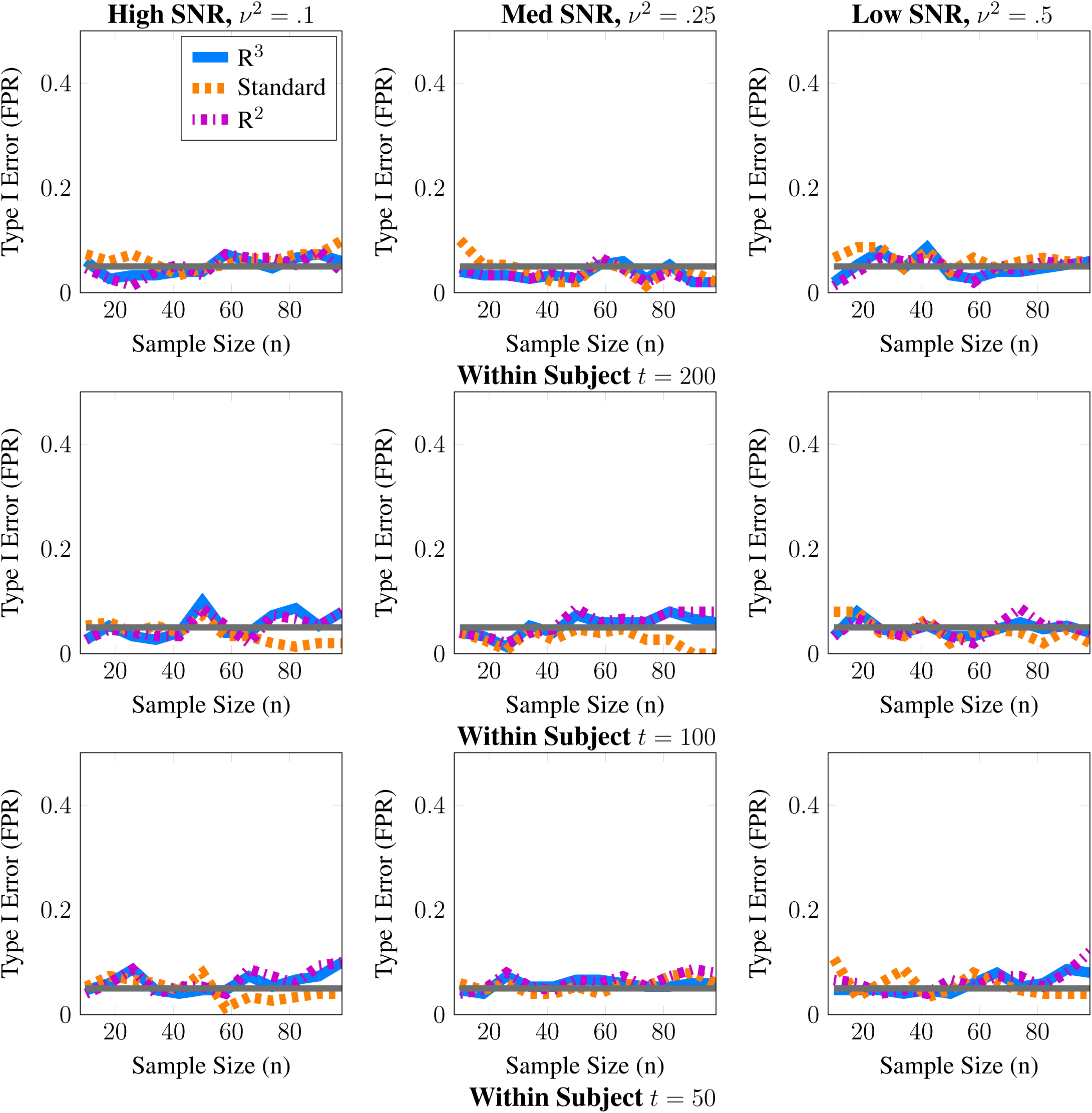
Statistical Type I Error Control for Node Density. These simulations evaluate the level of our tests; we report the estimated type-I error as a function of subject sample size *n*. The grey line represents the 5% level of the test. Here, we provide a complete set of Type-1 error simulations to complement the power analysis in Figure 3. All methods approximately control type I error across all scenarios studied for node density.

**Figure A.2.**
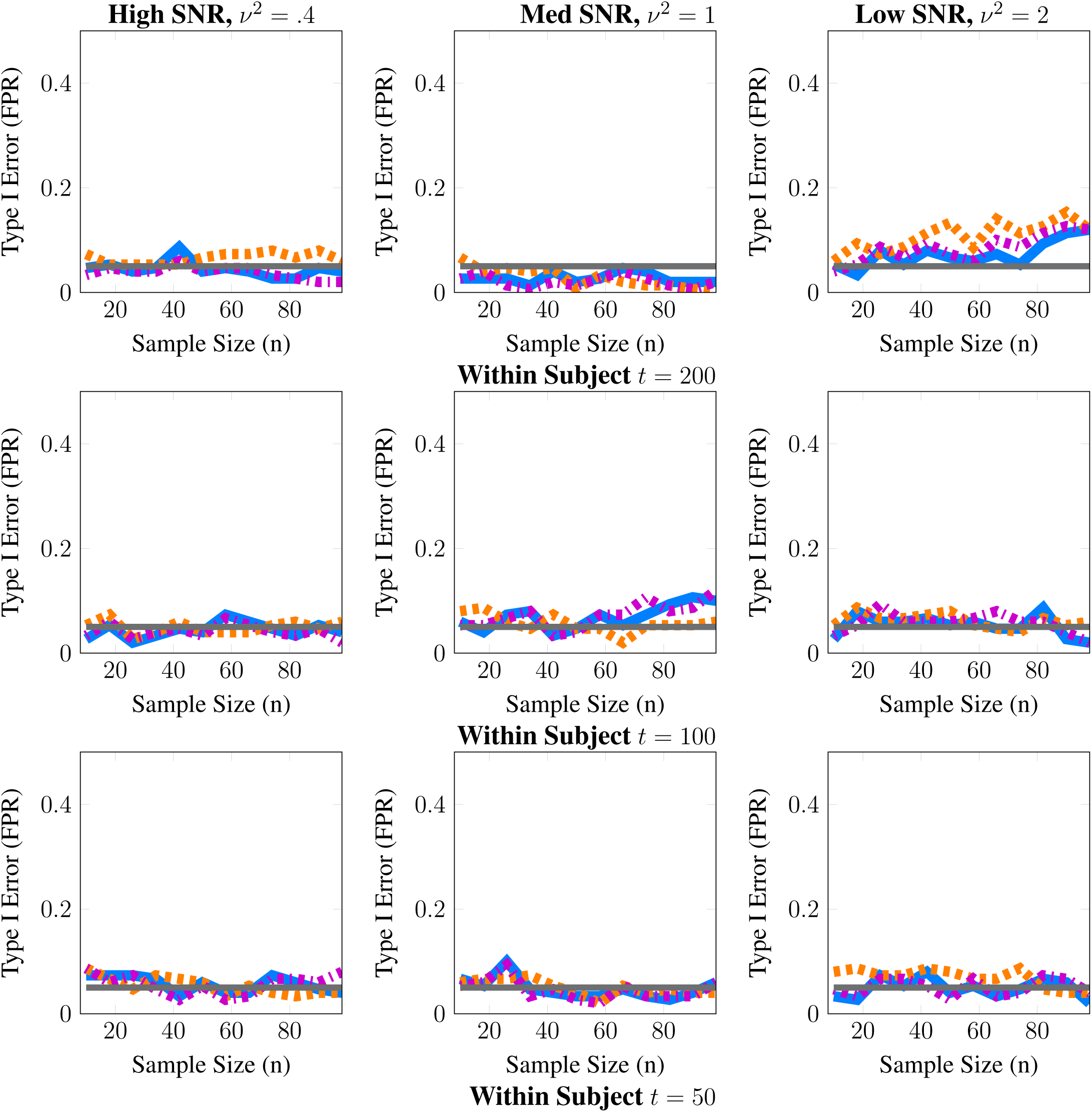
Statistical Type I Error Control for Subnetwork Density. These simulations evaluate the level of our tests; we report the estimated type-I error as a function of subject sample size *n*. The grey line represents the 5% level of the test. Here, we provide a complete set of Type-1 error simulations to complement the power analysis in Figure 4. All methods approximately control type I error across all scenarios studied for subnetwork density.

**Figure A.3.**
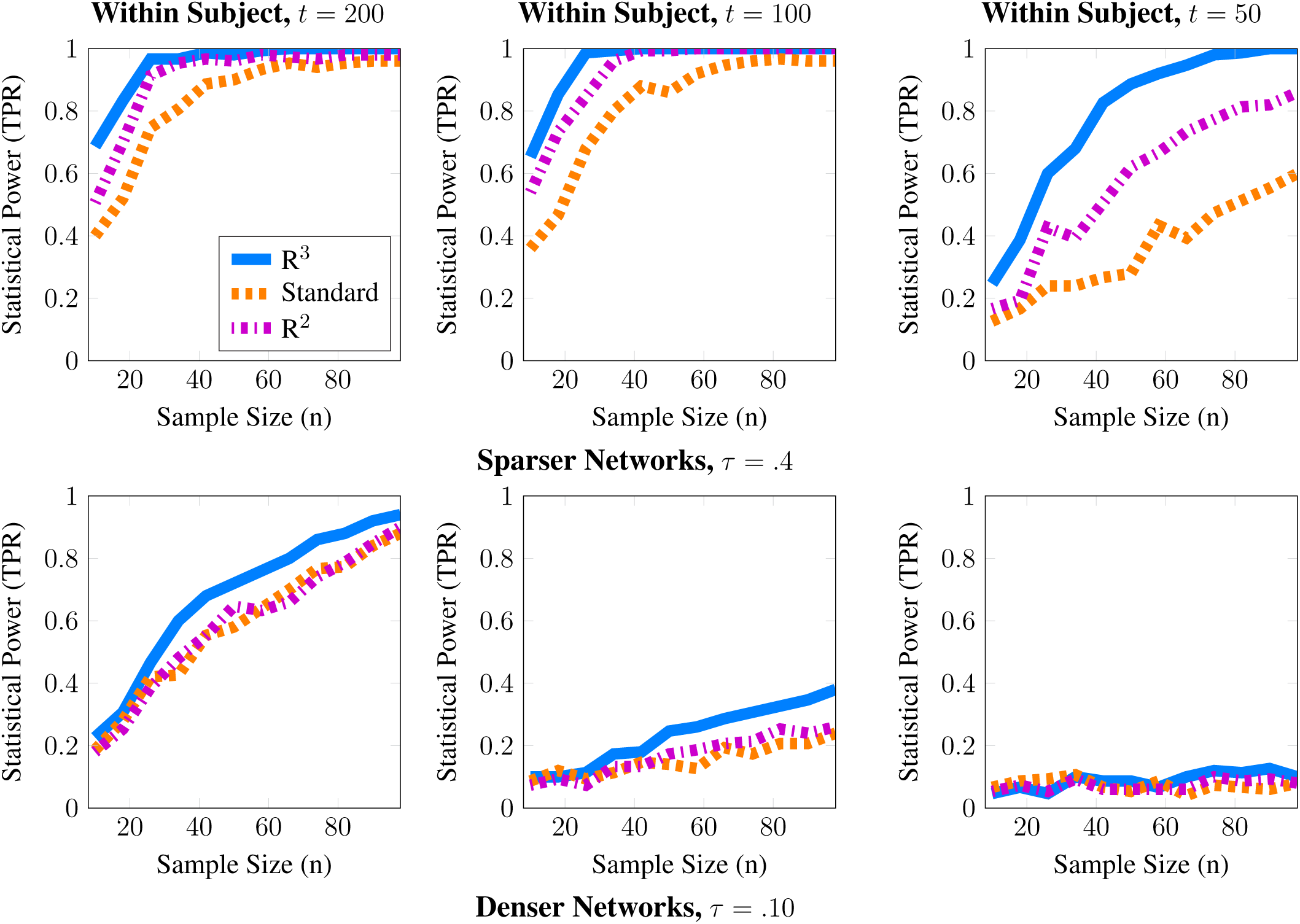
Statistical Power Analysis with Varying Baseline Sparsity. This figure complements the power analysis for node density in Figure 3 of the manuscript for the medium SNR case (*ν*^2^ = .25), where the number of nodes is *p* = 50. Whereas the baseline pseudo-real networks in Figure 3 consist of edges whose absolute partial correlation strength was greater than *τ* = .25, here we consider simulations where the baseline density is decreased (*τ* = .40) as well as increased (*τ* = .10). Notice that for the sparse baseline network, our results broadly match those of Figure 3. When node density varies with an explanatory covariate (*q* = 1), statistical power to detect this covariate effect improves with subject sample size *n* but crucially depends on the number of independent fMRI samples *t* from a single session. When networks are hard to estimate at limited within subject sample sizes *t* ≈ *p*, we expect estimates of node density to be both highly variable and potentially biased. However, as long as the baseline networks are sufficiently sparse, we can account for these errors via our methods **R**^3^ and **R**^2^. In fact, **R**^3^ achieves near perfect statistical power by adaptively improving the network metrics estimates of **R**^2^, thus improving statistical power over **R**^2^ and standard F-tests. In contrast, when baseline graphs are dense, and the sample sizes approach (*t* ≈ *p*), it becomes impossible to detect covariate effects. Thus, within subject sample sizes continue to be crucial for detecting covariate effects

The supplementary simulations in Figures A.1 & A.2 are consistent with Figure 5 of our manuscript, and demonstrate that all methods approximately control type-I error at the 5% level. In Figure A.3, as expected, statistical power decreases with smaller sample sizes, especially when *t* ≈ *p*. In the sparser baseline case, our methods, **R**^3^ and **R**^2^, are able to achieve better statistical power to detect covariate effects over standard F-tests. In the sparser network case, it is easier to estimate subject networks even in low sample sizes of *t* ≈ *p*, and initial stability scores continue to discriminate between true and false edges more effectively than in denser network regimes. Since the benefits of adaptive estimation depend on initial network estimates, we observe that the random adaptive penalization component of **R**^3^ improves the estimates of network metrics, thus achieving greater statistical power than **R**^2^ in sparser network regimes with small sample sizes. However, when baseline networks become denser, particularly when *τ* = .10, the ability of all methods to detect covariate effects begin to fail as within subject sample sizes reduce to *t* ≈ *p*. Overall our supplementary simulations continue to highlight the importance of within subject sample size *t*, and the benefits of our methods, **R**^3^ and **R**^2^ over the standard approach at smaller sample sizes.

### B TEST STATISTICS FOR **R**^3^ AND **R**^2^

R^3^ and **R**^2^ model resampled network metrics using repeated measures mixed effects models to account for two levels of variation in continuous network metrics. In this appendix, we begin with some elementary estimators and test statistics for covariate effects in the linear mixed effect (LME) model defined in Eq. (9) & (10) in Section 2.3 of our manuscript. Additionally, in Section B.2 we provide alternatives to the LME models in Section 2.3.3 of our manuscript for two levels of binary valued resampled statistics. As in the case of LME models, we outline relevant correlated binomial models and corresponding estimators for covariate effects.

#### B.1 ESTIMATORS FOR REPEATED MEASURES LME

Many estimators **Agresti** [2015] are available to estimate fixed effects for LME models, where the number of resamples within each subject is complete and balanced. We employ a generalization of ordinary least squares regression for correlated two-level data given by weighted least squares estimators (GLS) **Agresti** [2015]. Ideally, in order to make the least square residuals independent we weight the residuals by the precision matrix, 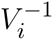, to obtain efficient estimates of *β*.

We redefine the earlier notation in Section 2.1 for the population model to account for the availability of resampled network metrics. We denote the overall design matrix by **W** = [*W*_1_ … *W*_*n*_]^⊤^. Here *W*_*i*_ is the *B* × (1 + *q* + *r*) subject level design matrix for the fixed effects, obtained by stacking centered and scaled explanatory and nuisance covariates [*X*_*i*_ *Z*_*i*_]. Let c denote a contrast vector to separate explanatory and nuisance covariates of interest such that *c* = [0 1_1×*q*_ 0_1×*r*_] and *c*^⊤^[*β* γ] = *β*_\0_. We omit the subscript excluding the intercept when referring to *β*_\0_ in this section. Here B denotes the number of resamples, *n* the number of subjects, q and r the number of explanatory and nuisance covariates, respectively.

Thus, the fixed effects estimate takes the form 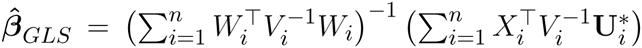 The corresponding partial Wald statistic for explanatory fixed effects is given by

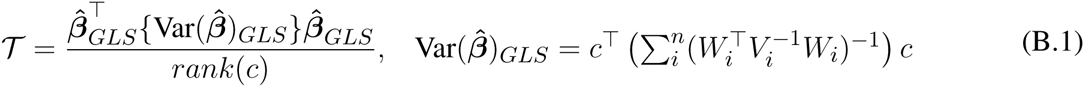

Since our two level model in Section 2.3.3 is a random intercept model for repeated measures, 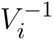 has compound symmetry structure and depends on two unknown parameters (*ν*^2^, *ϕ*^2^) that do not vary with subjects *i*. Consequently standard ANOVA and restricted maximum likelihood estimators for variance components, *ϕ*, *ν* coincide [**Searle et al.**, 2009], given by 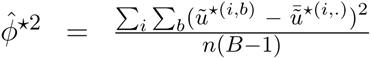 and 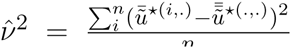 While Wald-type test statistics are asymptomtically *χ*^2^ distributed, they are better approximated by scaled F-distributions at finite samples. Finite sample corrections and estimates of the degrees of freedom for these F-distributions, provided by **Kenward and Roger** [1997], are widely adopted for inference in LME models to ensure better type-I error control. For more details on computational procedures and extensions to these models for more complex experimental designs, we refer the reader to **Agresti** [2015].

#### B.2 MIXED EFFECTS MODELS FOR CORRELATED BINARY DATA

As in the case of continuous metrics, when **R**^2^ and **R**^3^ produce resampled binary network statistics per subject, our data possesses two levels of variability. Although such statistics can be summarized using proportions 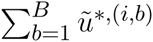 per subject, we cannot model these correlated proportions using binomial distributions, as the Îbinomial assumes all *nB* × 1 binary valued resampled statistics to be independent.

In fact, we expect the resampled statistics within each subject to be positively correlated. To resolve this problem, following the well established literature [**Liang and Hanfelt**, 1994; **Agresti**, 2015], we consider two-level models for correlated binary data.

To understand binomial models for correlated data, consider the example of the probability of observing an edge as the network metric of interest. Recall, from Eq.(6) that we seek to conduct inference over the fixed effect *β* which describes the rate of change in the subject edge probability in a population logit *π*_*i*_ = *η*_*i*_ = *Xβ* + *Z*_*γ*_ for a unit change in the covariate [**Williams**, 1982]. However we only observe network metrics for a sample of subjects in the population. To account for this inter-subject sampling variability, we introduce a continuous latent random variable *P*_*i*_ that takes values in the interval [0,1]. Additionally, however, we do not observe individual subject edge probabilities *P*_*i*_ but rather observe binary network statistics per subject. Thus, conditional on a subject’s true edge probability *P*_*i*_, we assume that each resampled network statistic *ũ*^*,(*i*,*b*)^ is Bernoulli distributed, such that *ũ*^*,(*i*,*b*)^|*P*_*i*_ = *p*_*i*_ ~ Ber(1,*p*_*i*_). Together, this gives us the following model for the observed proportions 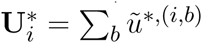

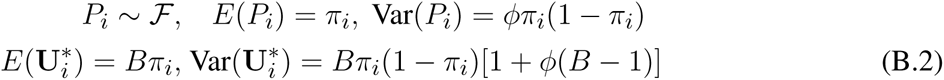

By employing this two-level model, we account for overdispersion in correlated resampled statistics in the form of the multiplicative correction term [1 +*ϕ*(*B* – 1)]. Note that, while we can specify a fully parametric model for 𝓕 using beta or correlated binomial distributions, specifying the first and second moments is adequate [**Williams**, 1982; **Searle et al.**, 2009] for the estimation and inference of fixed effects.

In the presence of balanced within subject resamples *B*, our two-level model (B.2) is very similar to our single level logistic-linear model in (6) with the exception of the additional overdispersion factor (1+*ϕ*(*B* – 1)). Thus, standard iterative reweighted least squares estimation can be used to obtain estimates of fixed effects *β*, γ and moment estimators for *ϕ* [**Kleinman**, 1973; **Williams**, 1982]. We proceed with inference for *β̂*, using Wald type statistics in (B.1), by ensuring that standard sample variance estimates for Var(*β*) incorporate the overdispersion factor. In the absence of balanced data, or for more complex experimental designs such as longitudinal imaging studies we recommend the maximum quasi-likelihood or generalized estimating equations [**Liang and Hanfelt**, 1994] for correlated binary data.

